# Periaxonal and nodal plasticity modulate action potential conduction in the adult mouse brain

**DOI:** 10.1101/726760

**Authors:** Carlie L Cullen, Renee E Pepper, Mackenzie T Clutterbuck, Kimberley A Pitman, Viola Oorschot, Loic Auderset, Alexander D Tang, Georg Ramm, Ben Emery, Jennifer Rodger, Renaud B Jolivet, Kaylene M Young

## Abstract

Central nervous system myelination increases action potential conduction velocity, however, it is unclear how myelination is coordinated to ensure the temporally precise arrival of action potentials, and facilitate information processing within cortical and associative circuits. Here, we show that mature myelin remains plastic in the adult mouse brain and can undergo subtle structural modifications to influence action potential arrival times. Repetitive transcranial magnetic stimulation and spatial learning, two stimuli that modify neuronal activity, alter the length of the nodes of Ranvier and the size of the periaxonal space within active brain regions. This change in the axon-glial configuration is independent of oligodendrogenesis and tunes conduction velocity to increase the synchronicity of action potential transit.

## Introduction

Within the central nervous system (CNS), oligodendrocytes elaborate myelin internodes to facilitate the rapid and saltatory conduction of action potentials and provide vital trophic support to axons via the periaxonal space (Simons and Nave, 2015). The speed of action potential conduction along individual axons is dependent on the molecular and structural parameters of the axon, including its diameter; the presence, length and thickness of myelin internodes; the length of the nodes of Ranvier; and the relative density of ion channels clustered at the nodes and paranodes (Arancibia-Carcamo *et al*., 2017; Ford *et al*., 2015; Freeman *et al*., 2016; Halter and Clark, 1993; Seidl, 2014; Young *et al*., 2013). Recently, it was also shown that axial conduction through the periaxonal space is important for the saltatory propagation of action potentials (Cohen *et al*., 2020). As even seemingly small changes in conduction velocity have the potential to significantly impact neuronal network function, by altering spike-time arrival or by disrupting the synchrony of brain wave rhythms (Pajevic *et al*., 2014), these structures must be precisely established and retain a degree of plasticity.

Oligodendrogenesis and myelination commence late in development (Jakovcevski *et al*., 2009; Kessaris *et al*., 2006; Lu *et al*., 2002) and peak prior to adolescence, however, new oligodendrocytes are generated throughout life (Dimou *et al*., 2008; Hill *et al*., 2018; Hughes *et al*., 2018; Rivers *et al*., 2008; Yeung *et al*., 2014; Young *et al*., 2013) to replace dying cells (Koenning *et al*., 2012; Yeung *et al*., 2014) and add myelin to previously unmyelinated or partially myelinated axons (Hill *et al*., 2018). The process of myelination is significantly influenced by experience, as social isolation early in life reduces the addition of myelin to the prefrontal cortex (Liu *et al*., 2012; Makinodan *et al*., 2012), and reducing visual input (monocular deprivation) shortens the myelin internodes elaborated by oligodendrocytes in the affected optic nerve (Etxeberria *et al*., 2016; Osanai *et al*., 2018). Conversely, in the adult brain, increasing neuronal activity, either by direct neuronal stimulation or through learning a new skill, promotes oligodendrogenesis and myelination of the activated circuits (Cullen *et al*., 2019; Gibson *et al*., 2014; Li *et al*., 2010; McKenzie *et al*., 2014; Mitew *et al*., 2018; Sampaio-Baptista *et al*., 2013).

In the developing zebrafish spinal cord, the extension and retraction of internodes is regulated by distinct patterns of calcium activity within the nascent myelin sheaths, and this can be partially regulated by neuronal activity (Baraban *et al*., 2018; Krasnow *et al*., 2018). In the mammalian brain, periodic and mitochondria-derived calcium transients can be detected within myelin sheaths that increase in frequency at the peak of cortical myelination and during remyelination in the adult mouse brain (Battefeld *et al*., 2019). Even after maturation, oligodendrocytes retain some capacity for internode remodeling, with a subset of internodes extending or retracting over time (Hill *et al*., 2018; Hughes *et al*., 2018). This capacity is perhaps best highlighted by the extension of established myelin sheaths to occupy an adjacent segment of recently demyelinated axon, following the ablation of a single myelinating oligodendrocyte in the zebrafish spinal cord (Auer *et al*., 2018). However, the extent to which mature oligodendrocytes remodel their internodes in the healthy brain, in response to specific physiological stimuli such as altered neuronal activity, has not been explored. Herein, we show that modulating neuronal activity, either artificially through repetitive transcranial magnetic stimulation (rTMS) or physiologically through learning, has no effect on gross internode length, but results in bi-directional adaptive changes to the axo-myelinic ultrastructure, that fine tune action potential conduction speed in the adult mouse brain.

## Results

### The gross myelinating morphology of mature oligodendrocytes does not change with iTBS

We have previously shown that non-invasive, low-intensity rTMS (Li-rTMS), delivered in an intermittent theta burst (iTBS) pattern, promotes the survival and maturation of new oligodendrocytes within the primary motor cortex (M1) (Cullen *et al*., 2019). To determine whether the non-invasive stimulation of M1 could induce adaptive changes in myelinating oligodendrocytes, we labelled a subset of mature cortical oligodendrocytes by giving a single dose of tamoxifen to adult (P83) *Plp-CreER :: Tau-mGFP* transgenic mice, one week prior to commencing Li-rTMS (see STAR methods). Following 14 days of sham-stimulation or iTBS we analyzed the morphology of mGFP-labelled myelinating oligodendrocytes within M1 (**Figure 1A-B**) and the underlying corpus callosum (CC; **Figure 1C-D**). More specifically, we measured the length of mGFP^+^ internodes that were flanked on each end by contactin-associated protein (CASPR)^+^ paranodes (**Figure 1C-D**). We found that internodes elaborated by pre-existing mGFP^+^ M1 oligodendrocytes were shorter (range ∼2-118μm, 30.28 ± 0.77 mean ± SD; **Figure 1E**, **F**) than those elaborated in the CC (range ∼10-104μm, 50.68 ± 1.02 mean ± SD; **Figure 1G**, **H**), but that iTBS did not alter the average length (**Figure 1F**, **H**) or length distribution (**Figure 1E**, **G**) of internodes in either region. The density of mGFP^+^ internodes within the CC (**Figure S1**) made it impossible to attribute internodes to any single cell within this region. However, we were able to determine the number of internodes maintained by individual oligodendrocytes within M1 and found that this was also unchanged by iTBS (sham: 29 ± 3; iTBS: 28 ± 2, mean ± SEM, t-test p=0.62, n=13 and 12 cells respectively), indicating that iTBS does not lead to detectable changes in the gross myelinating morphology of mature oligodendrocytes.

**Figure 1.**
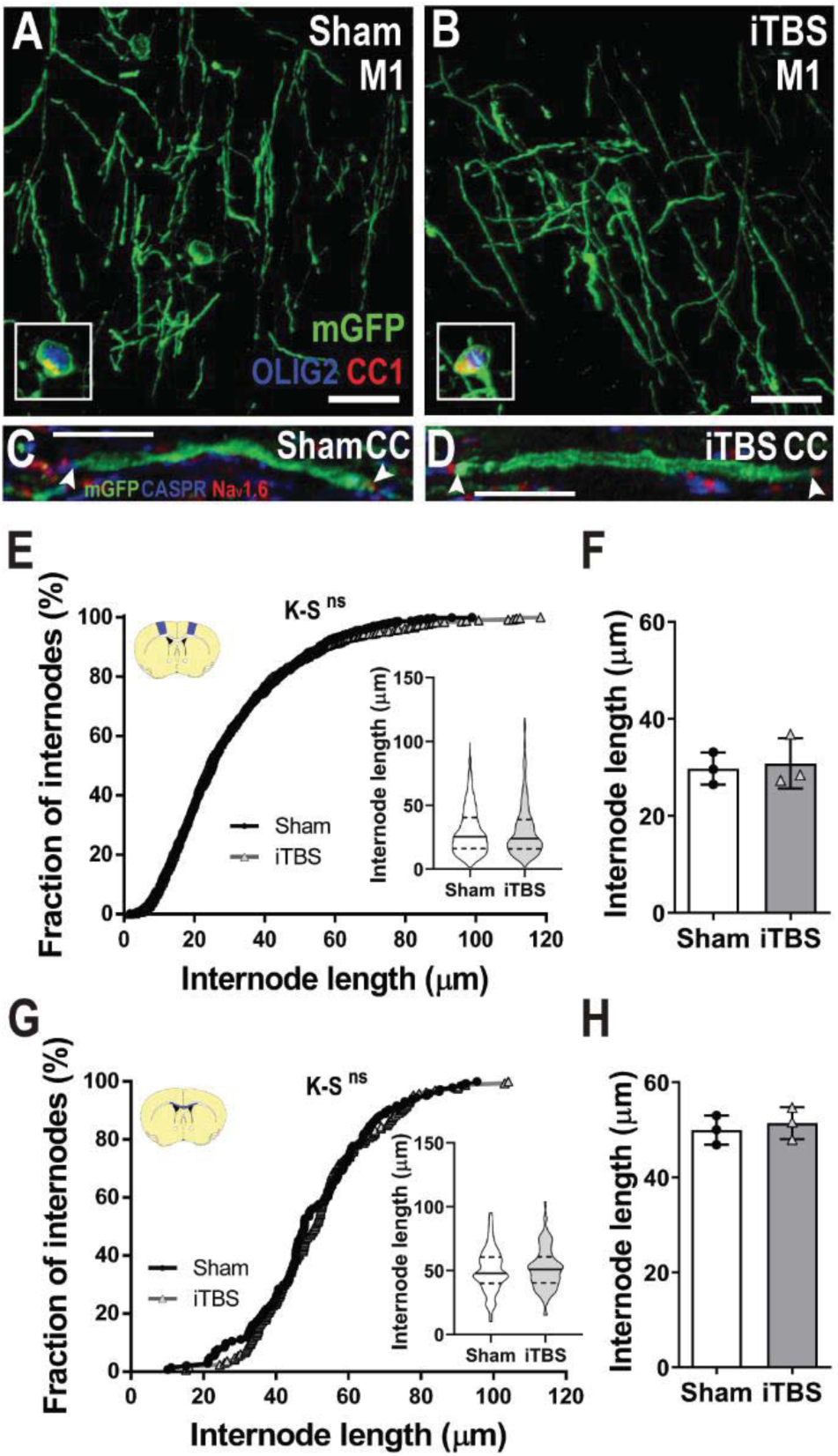
iTBS does not alter oligodendrocyte gross myelinating morphology. **A-B**) Representative compressed confocal z-stack of a mGFP^+^ (green) oligodendrocyte in the primary motor cortex (M1) of *Plp-CreER::Tau-mGFP* transgenic mice after 14 days of sham-stimulation (**A**) or iTBS (**B**). Inset, mGFP^+^ cell bodies show co-labeling for CC1 (red) and OLIG2 (blue). **C-D**) single mGFP^+^ internodes flanked on either end by CASPR^+^ paranodes (blue) and dense Na_v_1.6 staining at the node of Ranvier (red) in the corpus callosum (CC) of sham-stimulated (**C**) or iTBS (**D**) mice. **E-F**) Cumulative internode length distribution for mGFP^+^ internodes measured within M1 of sham (black circles) and iTBS (gray triangles) treated mice (E) [744 sham and 733 iTBS internodes; K-S test, K-S D=0.049, p=0.32; inset violin plot of internode length, Mann Whitney U (MWU) test p=0.63] and the average M1 internode length per animal (**F**) in sham (white bars) and iTBS (gray bars) treated mice [n=3 mice per group, t-test, t=0.31, p=0.77]. **G-H**) Cumulative internode length distribution for mGFP^+^ internodes measured within CC (**G)** [n=144 sham and n=166 iTBS internodes; K-S test, K-S D=0.08, p=0.63; inset violin plot of internode length, MWU test p=0.57] the average CC internode length per animal (**F**) sham and iTBS treated mice [n=3 mice per group, t-test, t=0.55, p=0.61]. Arrowheads indicate the end of an internode. Violin plots show the median (solid line) and interquartile range (dashed lines). Bars show mean ± SD. Scale bars represent 20μm.

### iTBS shortens nodes of Ranvier

At the end of each internode, anchoring proteins expressed by the myelin loops, such as neurofascin 155, interact with the axonal proteins contactin and CASPR to form paranodes (Bhat *et al*., 2001; Charles *et al*., 2002; Klingseisen *et al*., 2019; Peles *et al*., 1997; Sherman *et al*., 2005). These paranodal junctions maintain voltage gated sodium channels (Na_v_1.6) at the nodes of Ranvier (Freeman *et al*., 2016; Freeman *et al*., 2015; Suzuki *et al*., 2004). It has been suggested that node length is plastic, and may be modulated in response to altered neuronal activity to fine tune action potential propagation in accordance with information processing needs (Arancibia-Carcamo *et al*., 2017; Ford *et al*., 2015), however this has not yet been confirmed experimentally.

To determine whether iTBS affects specific axonal domains, we immunolabelled coronal brain sections from iTBS and sham stimulated animals to visualize paranodes (CASPR) and nodes of Ranvier (Na_v_1.6) (**Figure 2A-D**). By identifying regions of dense Na_v_1.6 staining that were clearly flanked by abutting CASPR^+^ paranodes, we quantified the length of individual nodes within M1 (**Figure 2A-B, E**) and the underlying CC (**Figure 2C-D, F**). We found that 14 days of iTBS treatment shifted node length distribution towards shorter nodes within both M1 (**Figure 2E**) and the CC (**Figure 2G**), which corresponded to a ∼19% reduction in average node length in M1 (**Figure 2F**) and a ∼16% decrease in the CC (**Figure 2H**). This shortening of the node length was not accompanied by a change in paranode length (or length distribution) (**Figure S2A-D**). To confirm the effect of iTBS on node length, we also performed high resolution stimulated emission depletion (STED) microscopy to visualize callosal nodes (Na_v_1.6) and paranodes (CASPR) in a separate cohort of animals following 14 days of iTBS or sham stimulation (**Figure S2E-F**). Consistent with our initial observations, we found that iTBS shifted the node length distribution towards shorter nodes (**Figure S2G**), corresponding to a reduction of ∼18% in average node length per mouse within the CC (**Figure S2H**).

**Figure 2.**
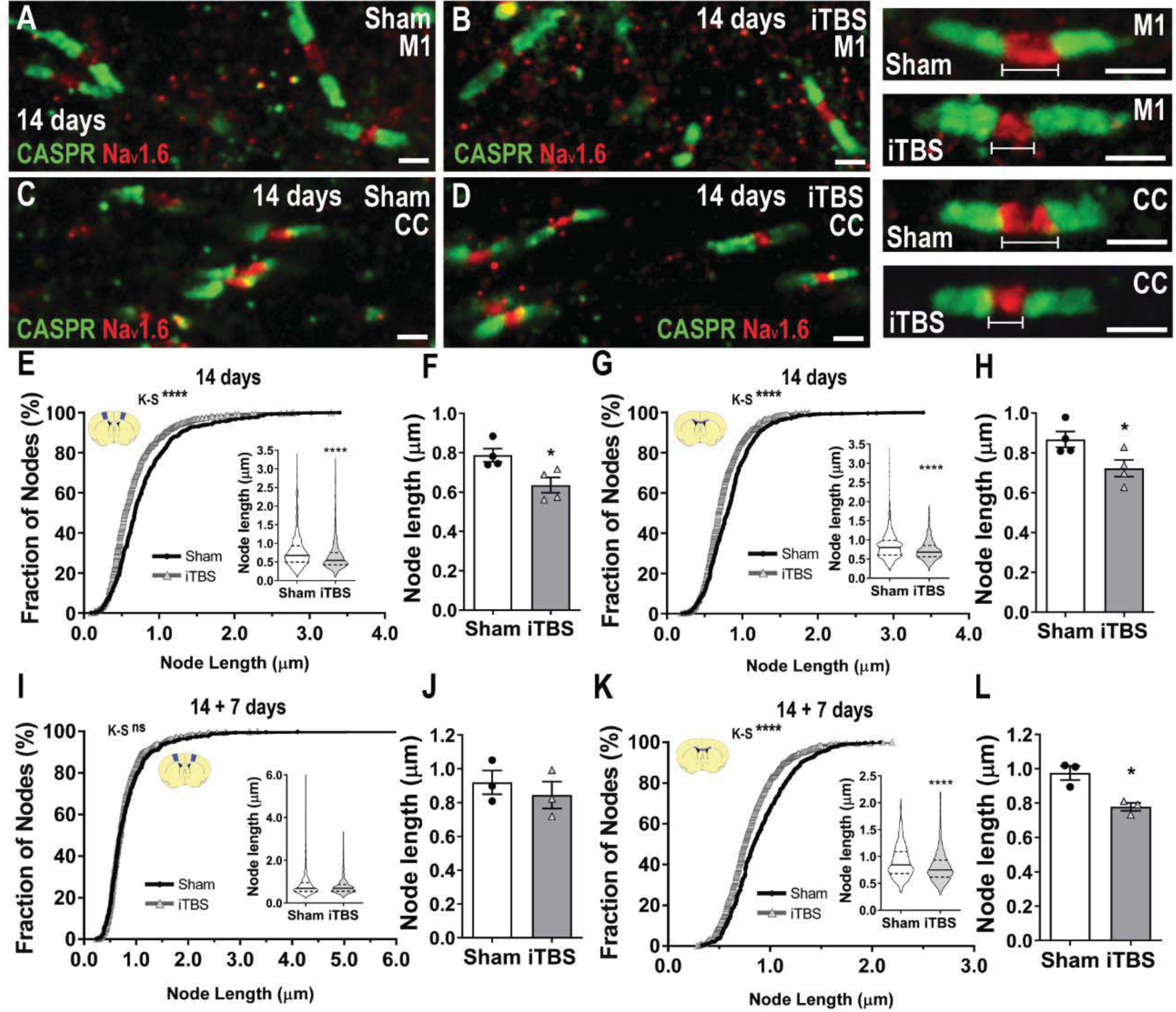
iTBS shortens nodes of Ranvier. **A-D**) Confocal images of nodes of Ranvier (Na_v_1.6; red) and paranodes (CASPR; green) in M1 (**A, B**) and the CC (**C, D**) after 14 days of sham-stimulation or iTBS. **E-F**) Cumulative node length distribution in M1 [656 sham (black circles) and 802 iTBS (gray triangles) nodes; Kolmogorov-Smirnov (K-S) test, K-S D=0.19, p<0.0001; inset, violin plot of node length, Mann Whitney U (MWU) test, p<0.0001] and average M1 node length per animal (**F**) in sham (white bars) and iTBS treated (gray bars) mice [n=4 mice per group, t-test, t=2.95, p=0.02]. **G-H**) Cumulative node length distribution in CC (**G)** [867 sham and 700 iTBS nodes; K-S D=0.13, p<0.0001; inset, violin plot of node length, MWU test, p<0.0001] and average CC node length per animal (**H**) in sham (white bars) or iTBS (gray bars) mice [n=4 mice per group, t-test, t=2.50, p=0.04]. **I-J**) Cumulative node length distribution in M1 (**I**) [452 sham and 576 iTBS nodes; K-S D=0.070, p=0.15; inset, violin plot of node length, MWU test, p=0.35] and average M1 node length per animal (**J**) in sham and iTBS treated mice [n=3 per group, t-test, t=0.70, p=0.52] 7 days after cessation of stimulation (14 + 7 days). **K-L**) Cumulative node length distribution in CC (**K**) [587 sham and 696 iTBS nodes; K-S D=0.16, p<0.0001; inset, violin plot of node length, MWU test, p<0.0001] and average CC node length per animal (**L**) in sham and iTBS treated mice [n=3 per group, t-test, t=4.2, p=0.01]. * p<0.05, ****p<0.0001. Violin plots show the median (solid line) and interquartile range (dashed lines). Bars show mean ± SD. Scale bars represent 1μm.

To determine whether node length was stable after stimulation ceased, we measured the length of individual nodes 7 days later (14+7 days). We found that nodes within M1 of iTBS mice returned to a similar average length and distribution as sham stimulated mice (**Figure 2I, J**), suggestive of bidirectional plasticity. However, within the CC the length distribution remained significantly shifted towards shorter nodes (**Figure 2K**), corresponding to a ∼20% decrease in average node length (**Figure 2L**), perhaps suggesting that nodal changes are more long-lived in the white matter relative to the grey matter.

### Spatial learning lengthens nodes of Ranvier

Learning physiologically modulates neuronal activity (Benchenane *et al*., 2010; Dupret *et al*., 2010; Dupret *et al*., 2013; Negrón-Oyarzo *et al*., 2018) and motor learning has been shown to promote oligodendrogenesis and increase myelination within the corpus callosum (Sampaio-Baptista *et al*., 2013; Xiao *et al*., 2016), but it is not clear whether mature oligodendrocytes respond to learning. Two weeks after tamoxifen administration (at P60), we trained adult (P74) *Plp-CreER :: Tau-mGFP* transgenic mice to learn the hippocampal dependent radial arm maze task (RAM; see STAR methods and **Figure S3A, C**). The subset of *Plp-CreER :: Tau-mGFP* transgenic mice that were exposed to the maze, but were not trained to learn the location of food rewards, are referred to as no-learning controls (**Figure S3B**). Following 14 days of no-learning (NL) or learning (L), we analyzed the morphology of mGFP-labelled myelinating oligodendrocytes within the hippocampal fimbria (**Figure 3A-B**), a major white matter tract that connects both hippocampi with subcortical and cortical regions including the thalamus and prefrontal cortex (Jin and Maren, 2015; Wyss *et al*., 1980). Due to the density of mGFP labelled internodes within the fimbria (**Figure S1C**), it was not possible to reliably attribute internodes to a single cell, however, by measuring the length of individual mGFP^+^ internodes (green), flanked by CASPR^+^ (blue) paranodes (**Figure 3C**), we determined that internodes within the fimbria were ∼5-109μm in length (**inset Figure 3D**), and that spatial learning had no effect on the average length or length distribution of internodes in this region (**Figure 3D-E**).

**Figure 3.**
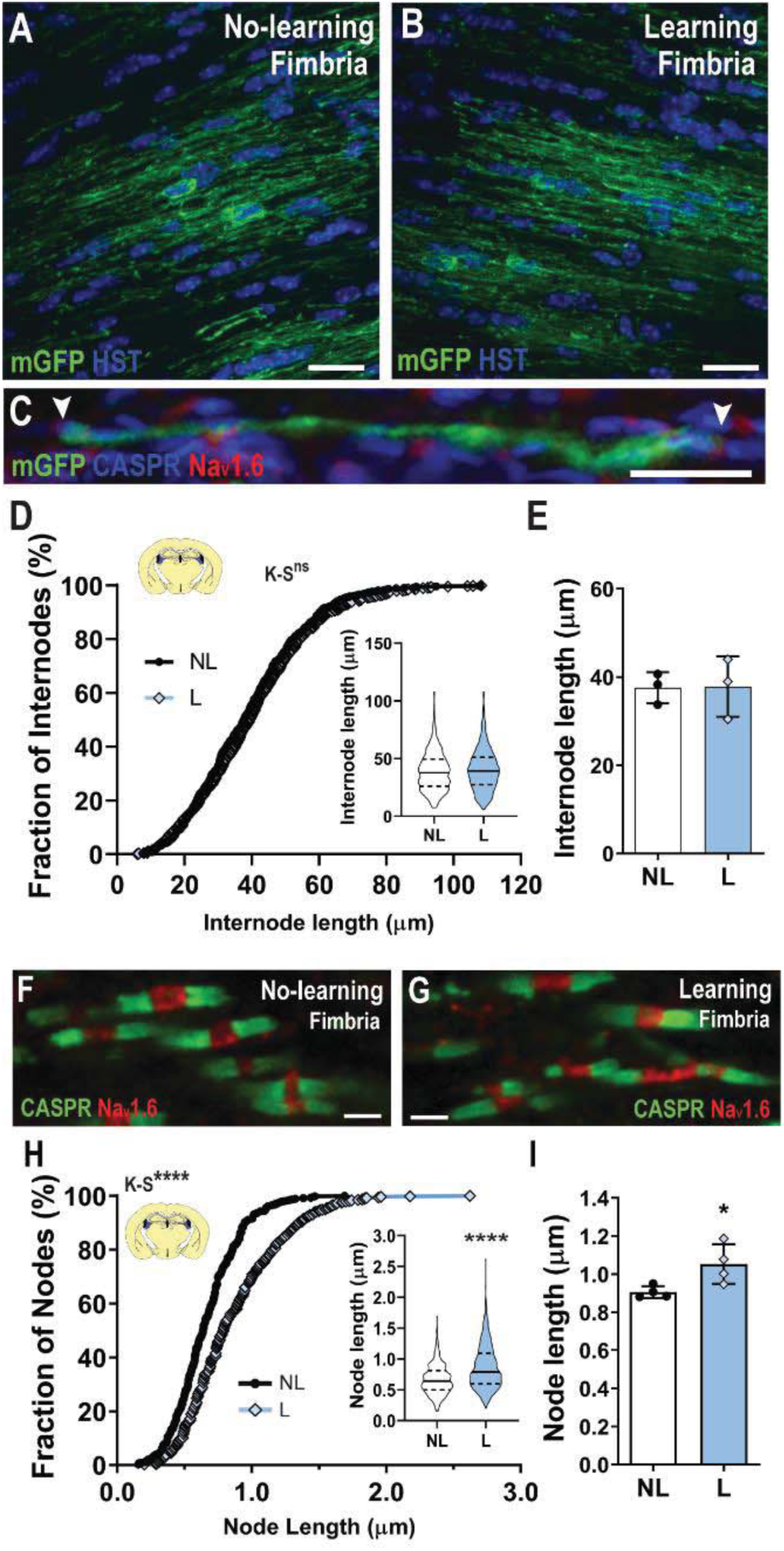
Spatial learning lengthens nodes of Ranvier. **A-B**) Representative compressed confocal z-stack of a mGFP^+^ (green) oligodendrocytes in the primary motor cortex (M1) of *Plp-CreER::Tau-mGFP* transgenic mice after 14 days of no-learning (**A**) or learning (**B**) in the radial arm maze. **C**) example of a single mGFP^+^ internode flanked on either end by CASPR^+^ paranodes (blue) and dense Na_v_1.6 staining (red) at the node of Ranvier. **D-E**) Cumulative internode length distribution for mGFP^+^ internodes measured within the fimbria of no-learning (NL; black circles) and learning (L; blue diamonds) mice (**D)** [n=532 NL and n=685 L; K-S test, K-S D=0.054, p=0.34; inset violin plot of internode length, Mann Whitney U (MWU) test p=0.15] and the average fimbria internode length per animal (**E**) in NL (white bars) and L (blue bars) mice [n=3 mice per group, t-test, t=0.053, p=0.96]. **F-G**) representative confocal images of nodes of Ranvier (Na_v_1.6; red) and paranodes (CASPR; green) in the fimbria of NL (**F**) and L (**G**) mice. **H-I**) Cumulative node length distribution in the fimbria (**H)** [448 NL and 520 L nodes; K-S D=0.25, p<0.0001; inset, violin plot of node length, MWU test, p<0.0001] and average fimbria node length per animal (**I**) in NL and L mice [n=4 per group, t-test, t=2.72, p=0.03]. * p<0.05, ****p<0.0001. Violin plots show the median (solid line) and interquartile range (dashed lines). Bars show mean ± SD. Scale bars represent 15μm (A-C), 1μm (F-G).

To determine whether learning induced nodal plasticity we quantified the length of nodes (Na_v_1.6, red) and paranodes (CASPR, green; **Figure 3F-G**) within the fimbria of no-learning and learning mice. Unlike iTBS, which shortened nodes, spatial learning produced a shift in node length distribution towards longer nodes (**Figure 3H**) and produced a corresponding ∼16% increase in average node length (**Figure 3I**). However, like iTBS, spatial learning did not alter the average length or length distribution of paranodes (**Figure S3D-E**). These data indicate that the nodes of Ranvier are dynamic and can lengthen or shorten in response to altered neuronal activity. Furthermore, these changes in node length occur without changing paranode length.

### Mature nodes of Ranvier are plastic

Oligodendrogenesis is driven by neuronal activity [reviewed by (Pepper *et al*., 2018)] and can influence node length (Schneider *et al*., 2016), making it possible that activity-induced nodal plasticity is a consequence of increased oligodendrocyte addition. However, this seems unlikely as oligodendrogenesis would be expected to consistently shorten nodes in the cortex and fimbria, and iTBS does not enhance oligodendrocyte addition to the CC (Cullen *et al*., 2019). To determine whether nodal plasticity can occur in the absence of oligodendrogenesis, tamoxifen was administered to P76 *Pdgfrα-CreER*^*TM*^ *:: Rosa26-YFP :: Myrf*^*fl/fl*^ (*Myrf*-deleted) mice, to fluorescently label OPCs and conditionally delete *myelin regulatory factor* (*Myrf*), a transcription factor essential for myelination and myelin maintenance (Emery *et al*., 2009). The conditional deletion of *Myrf* from adult OPCs reduced oligodendrocyte addition in M1 (p<0.05) and the CC (p<0.001) by >60% within 30 days of tamoxifen delivery (**Figure S4A-F**). When sham-stimulation or iTBS was initiated 14 days after Tx (P76+14), and the nodes of Ranvier (Na_v_1.6) imaged in the M1 at P76+28 (**Figure S4G-H**), iTBS again shifted the node length distribution towards shorter nodes (**Figure S4I**; K-S test, p<0.0001), and decreased average node length by ∼16% (**Figure S4J**; t-test p=0.04). Within the CC of *Myrf*-deleted mice, iTBS also shortened nodes (**Figure S4K**; K-S test, p<0.0001; **Figure S4L**; t-test, p=0.01), indicating that node shortening in response to iTBS cannot be entirely attributed to new oligodendrocyte addition.

To explore the possibility that nodal plasticity is instead mediated by mature, pre-existing oligodendrocytes, we selectively measured the length of already mature nodes of Ranvier (Na_v_1.6) following iTBS or sham-stimulation in *Plp-CreER :: Tau-mGFP* transgenic mice. These nodes were flanked by mGFP^+^ pre-existing internodes and formed prior to treatment (**Figure 4A-B**). Within M1, iTBS shortened the mature nodes (**Figure 4C**) and tended to decrease the average length of mature nodes in each mouse by ∼18% (**Figure 4D**, p=0.056). Within the CC, iTBS also shifted mature node length distribution towards shorter nodes (**Figure 4E**), corresponding to a ∼23% decrease in average node length (**Figure 4F**), and suggesting that node shortening is facilitated by mature oligodendrocytes.

**Figure 4.**
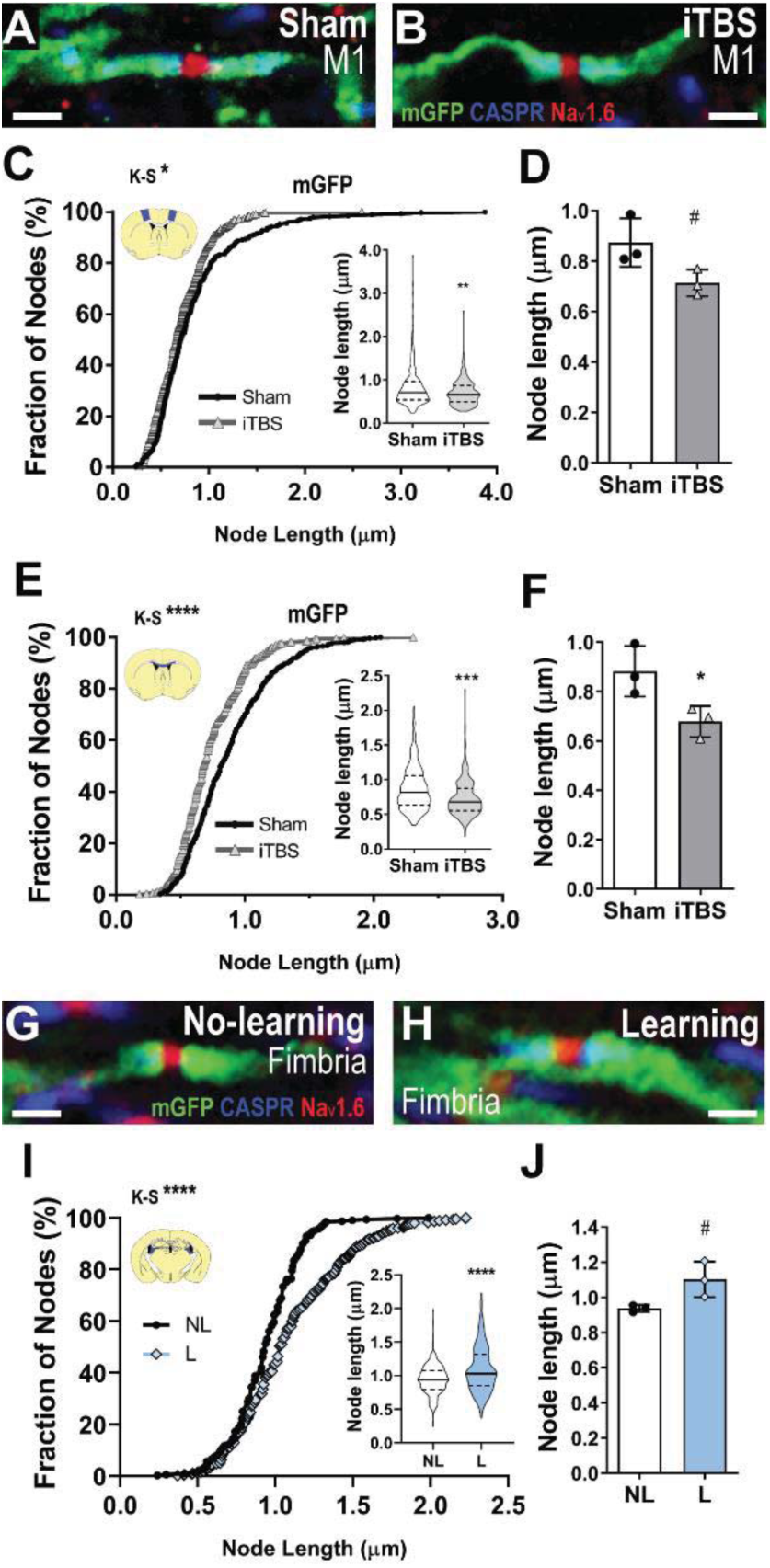
Nodal plasticity occurs between mature internodes. **A-B**) Mature node of Ranvier (Na_v_1.6; red) flanked by CASPR^+^ paranodes (blue) and mGFP^+^ internodes (green) in M1 of *Plp-CreER::Tau-mGFP* mice, formed prior to sham (**A**) or iTBS (**B**) treatment. **C-D**) Cumulative mature node length distribution in M1 (**C**) [273 sham (black circles) and 325 iTBS (gray triangles) mature nodes; K-S D=0.11, p=0.03; inset, violin plot of node length, MWU test, p=0.0019] and average mature node length per sham (white bars) and iTBS (gray bars) animal (**D**) [n=3 per group, t-test, t=2.52, p=0.056]. **E-F**) Cumulative mature node length distribution in the CC (**E)** [495 sham and 435 iTBS mature nodes; K-S D=0.22, p<0.0001; inset, violin plot of node length, MWU test, p<0.0001] and average mature node length per individual sham-stimulated and iTBS mice (**F**) [n=3 per group, t-test, t=2.94, p=0.04]. **G-H**) Mature node of Ranvier (Na_v_1.6; red) flanked by CASPR^+^ paranodes (blue) and mGFP^+^ internodes (green) in the fimbria of *Plp-CreER::Tau-mGFP* in mice that underwent no-learning (NL; **G**) or learning (L; **H**) in the radial arm maze. **I-J**) Cumulative mature node length distribution in the fimbria (**I**) [319 NL (black circles) and 453 L (blue diamonds) mature nodes; K-S D=0.25, p<0.0001; inset, violin plot of node length, MWU test, p<0.0001] and average mature node length per individual NL (white bars) and L (blue bars) mice (**J**) [n=3 per group, t-test, t=2.75, p=0.051]. ^#^ p<0.06, * p<0.05, ** p<0.01, *** p<0.001, **** p<0.0001. Violin plots show the median (solid line) and interquartile range (dashed lines). Bars show mean ± SD. Scale bars represent 1μm.

To determine whether node lengthening induced by spatial learning is also facilitated by mature oligodendrocytes, we similarly analysed nodes (Na_v_1.6) that were flanked by mGFP^+^ pre-existing internodes in the fimbria of no-learning (**Figure 4G**) and learning (**Figure 4H**) *Plp-CreER :: Tau-mGFP* transgenic mice. Consistent with our earlier observations, learning shifted mature node length distribution towards longer nodes (**Figure 4I**), which corresponded to an increase in the average length of mature nodes in each mouse of ∼17% (**Figure 4J**; p=0.051). These data suggest that mature oligodendrocytes can facilitate bi-directional, activity-induced nodal plasticity through the subtle expansion or retraction of existing myelin internodes.

### iTBS and spatial learning alter the size of the periaxonal space

For internodes to encroach on or retract from the nodes of Ranvier, the thickness or structure of myelin may be modified. For example, the sustained activation of extracellular signal-regulated kinases 1 and 2 (ERK1/2) in mature oligodendrocytes induces a marked decrease in node length in the CC of adult mice by increasing myelin thickness (Jeffries *et al*., 2016) and in the optic nerve, the thrombin-dependent detachment of paranodal loops led to thinner myelin (fewer myelin wraps) and longer nodes of Ranvier (Dutta *et al*., 2018). Therefore, we performed transmission electron microscopy (TEM) to quantify the g-ratio [axon diameter / (axon + myelin sheath diameter)] of myelinated axons in the CC of adult sham and iTBS treated mice (**Figure 5A-K**). We found that the distribution of g-ratios was left-shifted, indicating that iTBS treatment reduced axon g-ratio (**Figure 5A-B**), and this corresponded to a reduction of ∼7% in the average g-ratio of myelinated axons in iTBS mice compared with controls (**Figure 5C**).

**Figure 5.**
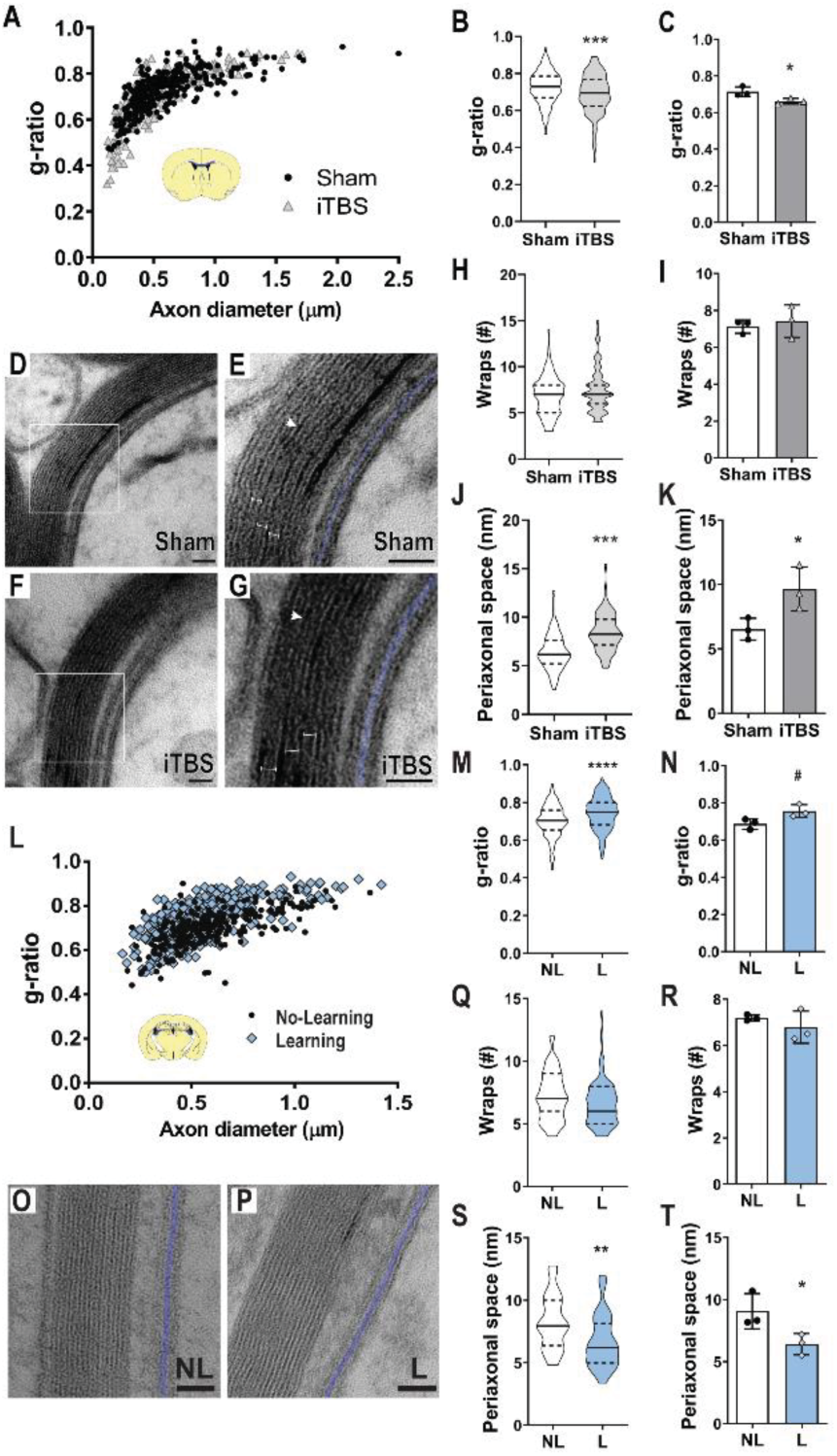
iTBS and spatial learning alter the size of the periaxonal space. **A**) Axonal diameter versus g-ratio for axons in the CC of sham-stimulated (black dots; n=323 axons) and iTBS (gray triangles; n=301 axons) mice [Kolmogorov-Smirnov (K-S) test for g-ratio D=0.18, p<0.0001]. **B**) Violin plot of g-ratio [323 sham (white) and 301 iTBS (gray) axons; Mann Whitney U (MWU) test p<0.0001]. **C**) Average g-ratio per individual sham and iTBS animal [t-test, t=2.98, p=0.04]. **D-G**) transmission electron microscopy (TEM) images of a myelinated axon in the CC of sham-stimulated (**D, E**) and iTBS (**F, G**) mice. White arrows = major dense line. Blue shading = periaxonal space. **H**) Violin plot of the number of myelin wraps per axon [104 sham and 125 iTBS axons; MWU test p=0.23]. **I**) Average number of myelin wraps per individual sham and iTBS animal [t-test, t=0.54, p=0.61]. **J**) Violin plot of periaxonal space width [75 sham and 60 iTBS axons, MWU test p<0.0001). K) Average periaxonal space width per individual sham and iTBS animal [t-test, t=2.84, p=0.04]. **L**) Axon diameter versus g-ratio for axons in the fimbria of no-learning (NL; black dots) or learning (L; blue diamonds) mice [209 NL and 374 L axons; K-S test for g-ratio, K-S D=0.24, p<0.0001]. **M)** Violin plot of g-ratio [209 NL axons (white) and 374 L axons (blue); MWU test, p<0.0001]. Average g-ratio per individual NL and L animal [t-test, t=2.64, p=0.057]. **O-P**) TEM images of myelinated axon in the fimbria of NL (**O**) and L (**P**) mice. **Q**) Violin plot of the number of myelin wraps per axon [55 NL and 55 L axons; MWU test, p=0.10]. **R**) Average number of myelin wraps per individual NL and L animal [t-test, t=0.98, p=0.38]. **S**) Violin plots of periaxonal space width [36 NL and 38 L axons; MWU test, p=0.0023]. **T**) Average periaxonal space width per individual NL and L animal [t-test, t=2.78, p=0.04]. Capped lines = single myelin wrap. ^#^ p<0.06, * p<0.05, ** p<0.01, *** p<0.001, **** p<0.0001. Violin plots show the median (solid line) and interquartile range (dashed lines). Bars show mean ± SD for n=3 animals per group. Scale bars = 25nm (D-G) or 50nm (O-P).

As axon diameter was equivalent between treatment groups (iTBS: 0.53 ± 0.17μm; sham: 0.58 ± 0.16μm, t-test p=0.09), the decreased g-ratio could reflect an increase in myelin thickness; however, when we quantified the number of myelin wraps around individual axons in the CC, by counting the major dense lines within each myelin sheath, we found that the number of myelin wraps was not affected by treatment (**Figure 5D-I**). Similarly, the average thickness of each wrap [the distance between each major dense line; capped lines in **Figure 5E, G**; sham: 8.3 ± 0.3nm, iTBS: 8.6 ± 0.2nm, mean ± SD, n=3 per group, t-test, t=1.27, p=0.27] and the width of the inner tongue process (sham: 30 ± 7nm; iTBS: 31.6 ± 3.2nm, mean ± SD, n=3 per group, t-test, t=0.35, p=0.73) were unchanged by iTBS treatment. Surprisingly, the width of the fluid-filled space under the internode, between the axon and the myelin sheath, known as the periaxonal space increased by ∼47% following iTBS (**Figure 5J-K**), perhaps suggesting that the myelin sheath is pushed outwards or reconfigured away from the axon.

Remarkably, spatial learning had the opposite effect on myelin ultrastructure in the fimbria (**Figure 5L-T**). Learning produced a significant leftward-shift in the g-ratio distribution for myelinated axons in the fimbria (**Figure 5L-M**), corresponding to a ∼9% increase in the average g-ratio (**Figure 5N**, p=0.057). This increase in g-ratio could not be explained by a change in axon diameter (no-learning: 0.61 ± 0.01μm, learning: 0.56 ± 0.11μm, mean ± SD; n=3 per group, t-test, t=0.87, p=0.42) or the number of myelin wraps elaborated (**Figure 5O-R**), but was associated with a ∼29% decrease in the average width of the periaxonal space (**Figure 5S-T**). These data suggest that bidirectional mechanisms link changes in neuronal activity with adaptive changes in myelin ultrastructure: iTBS is associated with an expansion of the periaxonal space and a contraction of the nodes of Ranvier for axons in the CC, while spatial learning is associated with a contraction of the periaxonal space and an extension of the nodes of Ranvier for axons in the fimbria.

### The size of the periaxonal space modulates conduction velocity

The frequency and composition of nodal domains can have a marked impact on action potential propagation along an axon (Ford *et al*., 2015; Freeman *et al*., 2016; Schneider *et al*., 2016), and node length is predicted to exert a strong influence on action potential conduction speed (Arancibia-Carcamo *et al*., 2017; Ford *et al*., 2015; Halter and Clark, 1993). Within the cortex, the estimated density of Na_v_1.6 channels at the node is directly proportional to node length, and this constant density of ion channels is predicted to induce a concave relationship between node length and speed of conduction, with very short or very long nodes causing a reduction in conduction velocity along axons (Arancibia-Carcamo *et al*., 2017). Furthermore, the inclusion or omission of a periaxonal space predicts whether new internodes, added to the mature brain, increase or decrease the simulated conduction velocity of an axon, respectively (Young *et al*., 2013). More recently, it was shown that electrical conductance within the periaxonal space facilitates the saltatory propagation of action potentials and that increasing the size of this space would theoretically slow conduction velocity (Cohen *et al*., 2020).

To determine how changes in node length and periaxonal ultrastructure might influence conduction speed, we performed computational simulations of action potential propagation in myelinated callosal axons by further adapting the mathematical model developed by Richardson and colleagues (Bakiri *et al*., 2011; Richardson *et al*., 2000) (**Figure 6A-B**; see STAR methods and **Figure S5**). Using this model, we initially explored how the size of the periaxonal space might impact conduction velocity (**Figure 6C-D**). In line with Cohen et al. (2020), we find that simulated conduction velocity is up to 3.5 times slower when the periaxonal space width is set to 20nm rather than 0nm (**Figure 6C**), which equates to a conduction delay of up to ∼9ms (at 21°C) over a distance of 1cm (**Figure 6D**).

**Figure 6.**
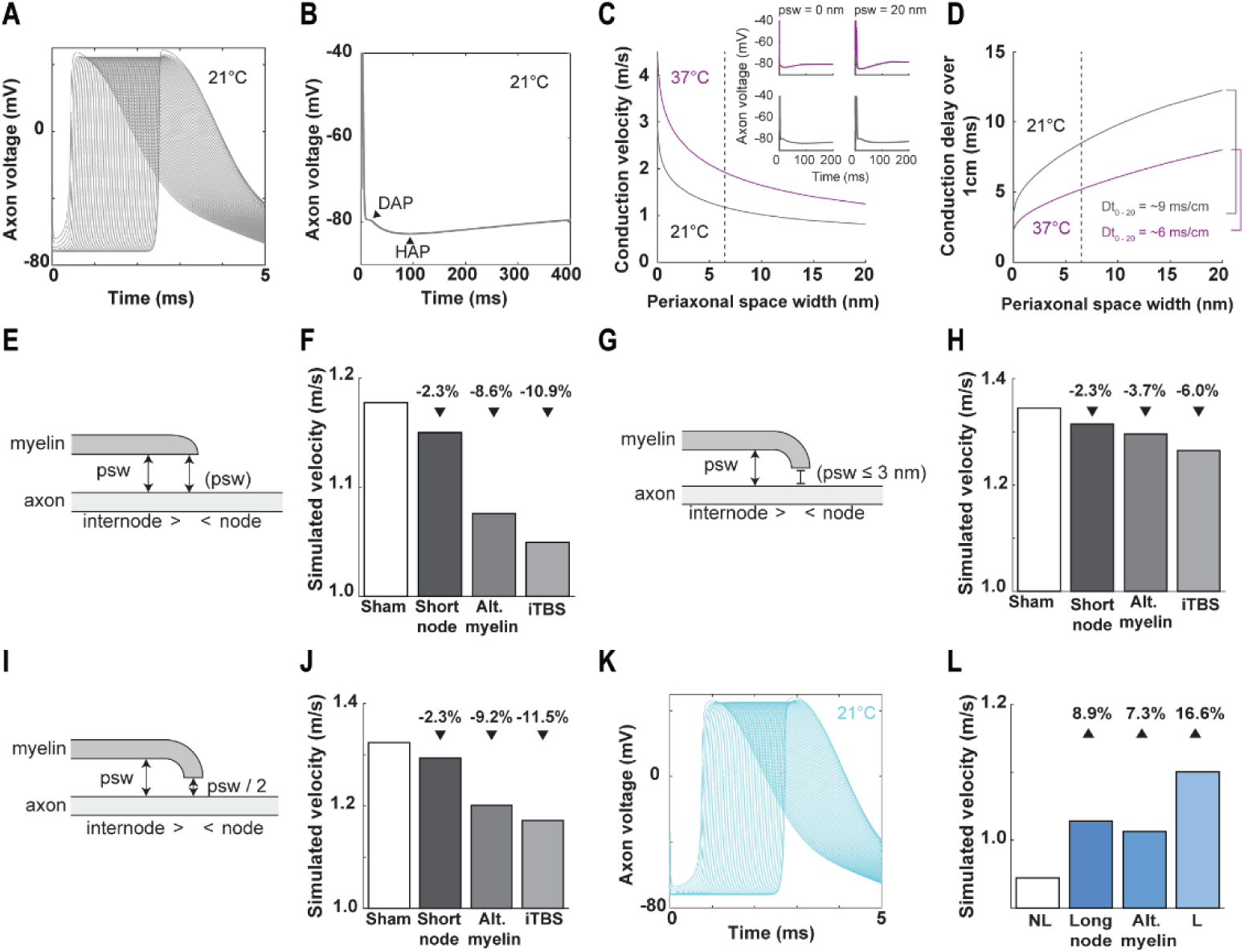
The size of the periaxonal space modulates conduction velocity. **A**) Action potentials simulated at consecutive callosal nodes at 21°C. **B**) The extended time-course of action potentials generated by the model at 21°C, highlighting a slight depolarizing after-potential (DAP) and a hyperpolarizing after-potential (HAP). **C**) Simulated conduction velocity of callosal axons relative to periaxonal space width (*psw*) at 37°C (magenta) and 21°C (gray), dashed line indicates average *psw* following sham stimulation insets show action potential waveforms at the extremities of the tested range (*psw* = 0 or 20nm) at 21°C and 37°C. **D**) Conduction delay over 1cm relative to *psw* at 21°C and 37°C. **E**) Schematic showing model parameters in which *psw* at the paranode is uniform to internode *psw*. **F**) Predicted conduction velocity of a sham stimulated (white) axon versus an axon with either the node length shortened, the periaxonal space widened or both (iTBS) using the model depicted in (E). **G**) Schematic showing model parameters in which *psw* at the paranode is set to ≤ 3nm. **H**) Predicted conduction velocity of a sham stimulated axon versus an axon with either the node length shortened, periaxonal space widened or both (iTBS) using the model depicted in (G). **I**) Schematic showing model parameters in which *psw* at the paranode is set to half internode *psw*. **J**) Predicted conduction velocity of a sham stimulated axon versus an axon with either the node length shortened, periaxonal space widened or both (iTBS) using the model depicted in (I). **K**) Action potentials simulated at consecutive nodes within the fimbria at 21°C. **L**) Predicted conduction velocity of a no-learning (NL) control axon within the fimbria (white) versus an axon with either longer nodes, a narrower periaxonal space or both (learning, L).

By initially configuring the model parameters so that the periaxonal space was uniform under the internode and paranode (**Figure 6E**), and setting all other parameters to match those measured under sham conditions (**Figure 6F; Table S1**), we obtained a theoretical conduction velocity of 1.18m/s at 21°C (1.91m/s at 37°C; **Figure S5**). Reducing the node length, as occurred during iTBS (**Figure 2; Table S1**), slowed action potential propagation by ∼2.3%, while altering the myelin sheath parameters (decreased g-ratio + increased periaxonal space; **Figure 5; Table S1**) exerted a greater influence on conduction speed, effectively slowing propagation by ∼8.6% (**Figure 6F**). Implementing all of the ultrastructural changes observed following iTBS (reduced node length and g-ratio + increased periaxonal space; **Figure 2, Figure 5, Table S1**) had an additive effect, slowing conduction velocity by ∼10.9% (to 1.05m/s; **Figure 6F**). This effect was exaggerated at 37°C with iTBS inducing a theoretical ∼12.3% reduction in conduction speed (**Figure S5D**).

We then changed the model parameters so that the periaxonal space under the internode could change, while the periaxonal space at the paranode did not exceed 3nm (**Figure 6G**), which is the middle of the size range that has been reported elsewhere (Nans *et al*., 2011; Rosenbluth, 1995; Waxman *et al*., 1995). In this scenario, the simulated conduction velocity for sham-stimulated axons was slightly faster, being 1.34m/s at 21°C (**Figure 6H;** 37°C data shown in **Figure S5F**). Reducing node length still slowed conduction velocity by ∼2.3%, but the effect of altering the myelin sheath parameters was diminished, slowing conduction by only ∼3.7% and the combined effect of reduced node length + altered myelin (iTBS) was also weaker, inducing a theoretical ∼6% reduction in conduction speed (**Figure 6H**) which was similar at 37°C (∼5.7% reduction, **Figure S5F**). In a third model, we set the periaxonal space width at the paranode to be half of the width of the periaxonal space under the internode, creating a scenario in which the periaxonal space is narrower at the paranode, but changes proportionally with the internodal periaxonal space (**Figure 6I**). In this scenario, the simulated conduction velocity using sham parameters was 1.32m/s (**Figure 6J**) at 21°C (37°C data shown in **Figure S5H**), but the reduction in conduction velocity induced by altering myelin sheath parameters (∼9.2%) or by mimicking the iTBS condition (∼11.5%, **Figure 6J**; 12.5% at 37°C, **Figure S5H**) was comparable to the case where periaxonal space width was homogenous along the internode and paranode (**Figure 6E-F**).

Finally, we modelled action potential conduction in the fimbria using a parameter set matching the experimental observations obtained in the no-learning condition (**Figure 6K-L**; **Table S1**). The simulated conduction velocity along fimbria axons in the no-learning (NL) condition was 0.95m/s (**Figure 6L**) at 21°C (37°C data shown in **Figure S5L**), which is far slower than the simulated velocity in the CC, but is within range of measured conduction velocities for this region (Corcoba *et al*., 2015; Jones *et al*., 1999). Increasing the node length by ∼30%, to reflect the change produced by RAM learning (**Figure 3**), increased the simulated conduction speed by 8.9%, while altering the myelin sheath parameters (increased g-ratio + decreased periaxonal space, **Figure 5, Table S1**) increased conduction velocity by 7.3%. Again, implementing all of the changes measured following learning (L; increased node length and g-ratio + decreased periaxonal space; **Figure 3, Figure 5, Table S1**) had an additive effect, increasing conduction velocity by 16.6% (**Figure 6L**). At 37°C, this would be predicted to correspond to a 21.6% increase in conduction velocity following learning (**Figure S5L**). These data suggest that adaptive structural changes at nodes of Ranvier and the periaxonal space act in concert to slow down or speed up action potential conduction along an axon.

### iTBS slows conduction velocity across the CC

As our model predicted that iTBS would, on average, decrease action potential conduction velocity along axons in the CC, we decided to measure this by performing *ex vivo* field potential recordings of compound action potentials (CAPs) in the CC of sham and iTBS mice (**Figure 7A**). We found that the average conduction velocity along myelinated (M) CC axons in sham stimulated mice was 1.27 ± 0.19 m/s (**Figure 7B**), which is remarkably similar to the speed predicted by our simulations (**Figure 6J**). Conduction velocity along myelinated axons was reduced by ∼18% following iTBS (1.04 ± 0.19 m/s; **Figure 7B**). The amplitude of the myelinated axon component of the CAP also increased by ∼40% following iTBS (**Figure 7C**), and the half-width decreased by ∼9% (**Figure 7D**), suggesting that a greater number of action potentials arrived simultaneously at the recording electrode. The unmyelinated axon (UM) component of the CAP was not altered by iTBS treatment (**Figure 7A-D**). These data validate our *in silico* predictions and suggest that oligodendrocytes can make subtle changes to their myelin ultrastructure that impact the length of nodes of Ranvier and the size of the periaxonal space, and this allows them to fine-tune conduction velocity and enhance the coordinated conduction of action potentials within the active circuit.

**Figure 7.**
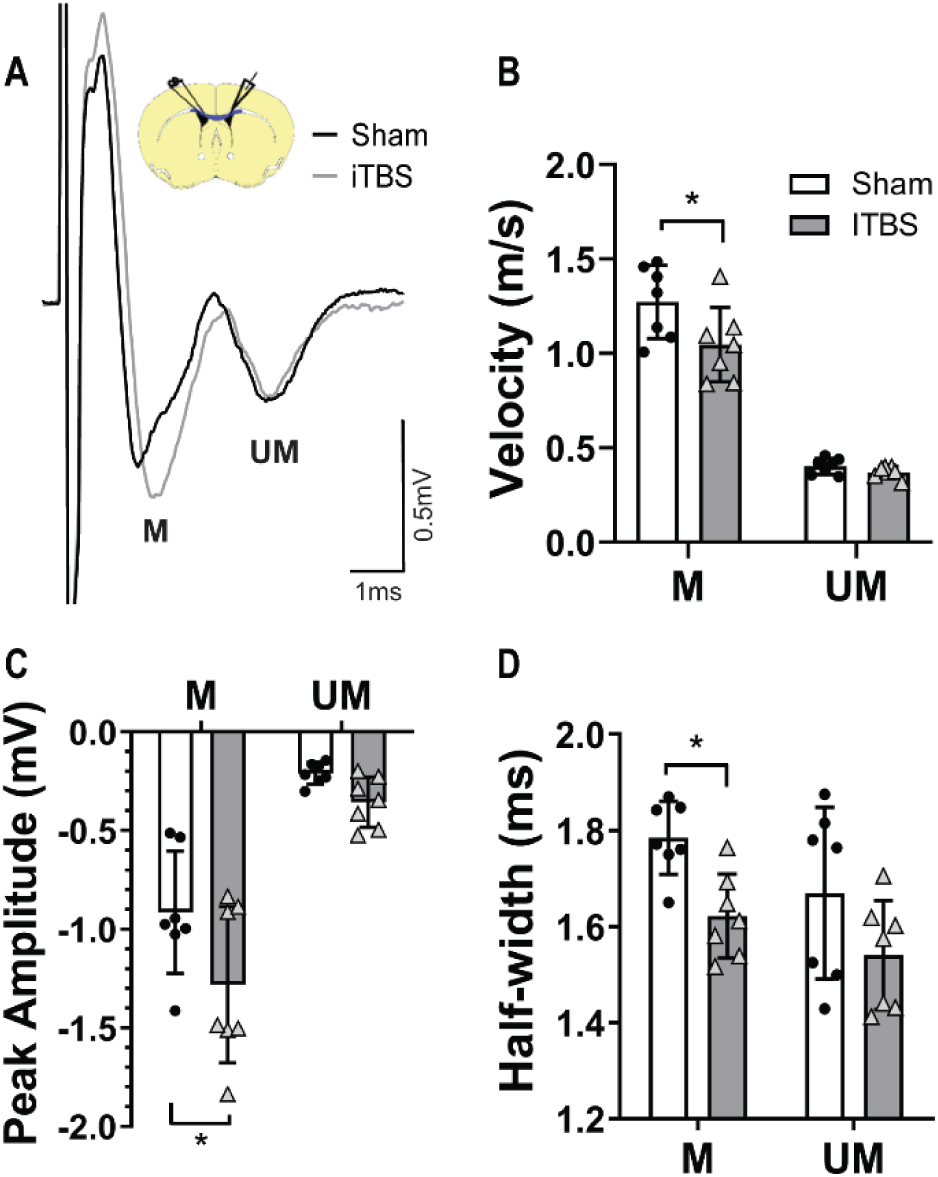
iTBS slows conduction velocity. **A**) Compound action potential (CAP) recorded in the CC of sham-stimulation (black) or iTBS (gray) mice. **B**) Conduction velocity of myelinated (M) and unmyelinated (UM) axons in sham-stimulated (black dots) or iTBS (gray triangles) mice [2-way ANOVA: interaction F(1,24)=3.37, p=0.078; axon population F(1,24)=211.1, p<0.0001; treatment F(1,24)=5.81, p=0.023]. **C**) Peak amplitude of M and UM axons [2-way ANOVA: interaction F(1,24)=1.25, p=0.27; axon population F(1,24)=68.10, p<0.0001; treatment F(1,24)=6.75, p=0.015]. **D**) Half-width of CAP peak corresponding to M and UM axons [2-way ANOVA: interaction F(1,24)=0.13, p=0.71; axon population F(1,24)=4.63, p=0.04; treatment F(1,24)=10.22, p=0.003]. Bars show mean ± SD. * p<0.05 by Bonferroni’s post-test.

## Discussion

### Are mature oligodendrocytes adaptable?

In recent years, it has become increasingly evident that central myelination is adaptive [see (Almeida and Lyons, 2017; Bechler *et al*., 2017; Mount and Monje, 2017) for reviews] and that changing the pattern of myelination within a circuit could have a profound impact on neural processing (Pajevic *et al*., 2014). This area of research has largely focussed on the addition of new myelin over time and in response to altered neuronal activity (Cullen *et al*., 2019; Gibson *et al*., 2014; Li *et al*., 2010; McKenzie *et al*., 2014; Mitew *et al*., 2018; Sampaio-Baptista *et al*., 2013; Xiao *et al*., 2016; Young *et al*., 2013), but we do not yet know whether existing oligodendrocytes adapt to altered neuronal activity and whether this affects the propagation of action potentials. Using a combination of fluorescence microscopy, ultrastructural analysis, computational modelling and electrophysiological recordings, we now show that the nodes of Ranvier and the periaxonal space undergo bi-directional plasticity in response to the artificial or physiological modulation of neuronal activity and that this response is likely mediated by mature oligodendrocytes. Furthermore, these adaptive changes in axo-myelin ultrastructure synergistically fine-tune conduction velocity along myelinated axons within the active circuit.

Once formed, mature myelinating oligodendrocytes are long lived (Tripathi *et al*., 2017) and the majority of the internodes they support are stable over time, however, there is some evidence that these mature cells retain the capacity to adjust internode length if required (Auer *et al*., 2018; Hill *et al*., 2018). We sought to determine whether changes in neural circuit activity could trigger adaptive changes in mature internodes but found that neither iTBS or spatial learning induced an overt extension or retraction of mature internodes (**Figures 1 & 3**). These data are consistent with fluorescent live imaging data showing that existing myelin sheaths within the somatosensory cortex do not change length in response to sensory enrichment or deprivation (Hughes *et al*., 2018). However, we observed marked alterations in the length of the nodes of Ranvier following both iTBS and spatial learning. Low intensity iTBS reduced node length within M1 and the underlying CC (**Figures 2, 4, S2 & S4**), while spatial learning increased node length within the hippocampal fimbria (**Figures 3 & 4**). These dynamic changes appear to be mediated by subtle adaptations made by mature oligodendrocytes, as preventing new oligodendrocyte addition did not impact iTBS-induced node shortening (**Figure S4**), and both iTBS and learning could alter the length of nodes that were already mature (flanked by pre-existing myelin internodes) prior to the change in activity (**Figure 4**). Previous studies have reported pathological node lengthening (Hinman *et al*., 2015; Howell *et al*., 2006; O’Hare Doig *et al*., 2017; Reimer *et al*., 2011) associated with myelin loss (Howell *et al*., 2006) or paranodal pathology (Huff *et al*., 2011; Reimer *et al*., 2011), however, we found no evidence of myelin pathology following iTBS or learning, which is a natural, physiological stimuli. Others have suggested that node length may be plastic and physiologically modulated (Arancibia-Carcamo *et al*., 2017; Ford *et al*., 2015) but, to our knowledge, this is the first experimental evidence of activity-induced adaptive plasticity at the node of Ranvier *in vivo*.

### Plasticity of the periaxonal space

iTBS decreased the g-ratio of myelinated axons in the CC. This is consistent with various other forms of cortical activity modulation, including the direct optogenetic stimulation of layer V pyramidal neurons in the premotor cortex (Gibson *et al*., 2014), the pharmacogenetic stimulation of layer 2/3 pyramidal neurons in the primary somatosensory cortex (Mitew *et al*., 2018) and the optogenetic inhibition of parvalbumin^+^ interneurons in the anterior cingulate cortex (Piscopo *et al*., 2018), that similarly decreased the g-ratio of axons within the underlying CC. However, we found that the decrease in g-ratio observed following iTBS was not the result of increased myelin thickness, or decreased axon diameter, but was instead associated with a marked increase in the periaxonal space (**Figure 5**). Surprisingly, spatial learning instead increased the g-ratio of myelinated axons within the hippocampal fimbria and was this was associated with a decrease in the size of the periaxonal space (**Figure 5**). This is the first report of an ultrastructural adaptation of the periaxonal space in response to altered neuronal activity. However, the periaxonal space has been reported to undergo pathological changes. For example, the periaxonal space is diminished in mice deficient in myelin associated glycoprotein (MAG) (Georgiou *et al*., 2004; Li *et al*., 1994; Quarles, 2007) – a protein expressed specifically in the periaxonal myelin membrane (Pronker *et al*., 2016), and vacuoles form within this space along optic nerve axons following connexin 32 / 47 knock-out (Menichella *et al*., 2006). Marked pathological swelling of the periaxonal space has also been observed following acute ischemic white matter injury and demyelination (Aboul-Enein *et al*., 2003). However, vacuolisation, or an uneven separation of the myelin from the axons was not observed in our tissue, suggesting that iTBS- or learning-induced modulation of the periaxonal space is not pathological, but a physiological adaptation to altered neuronal activity.

### Nodal and periaxonal plasticity modulate action potential conduction

The periaxonal space has been hypothesised to act as a receptacle for calcium, glutamate and potassium released by the axon during action potential propagation, as well as oligodendrocyte derived lactate and pyruvate, which is then shuttled through monocarboxylate transporters to provide metabolic support to the axon [reviewed in (Micu *et al*., 2017)], and as an important regulatory element of action potential conduction. The inclusion or omission of a periaxonal space predicts whether the addition of new internodes to an axon increased or decreased simulated conduction velocity along that axon (Young *et al*., 2013) and experimentally-constrained cable modelling revealed that the periaxonal space is almost certainly conductive and that current flow through this fluid filled space helps facilitate the saltatory propagation of action potentials along myelinated axons (Cohen *et al*., 2020). In line with our own data (**Figure 6**), these authors also show a marked theoretical slowing of conduction speed as the width of the periaxonal space is increased (Cohen *et al*., 2020).

Previous ultrastructural analyses of myelinated axons in the CNS have shown that the periaxonal space is typically narrower at the paranode than under the internode (Nans *et al*., 2011; Rosenbluth, 1995; Waxman *et al*., 1995) and the capacity for ultrastructural adaptation of the periaxonal space to influence axonal conduction speed is likely to be dependent on its size both under the internode, and at the paranode. As we were unable to measure periaxonal space width at the paranode, we instead modelled action potential propagation along callosal axons following sham or iTBS treatment, incorporating three possible paranodal scenarios: i) that the periaxonal space width was uniform under the internode and paranode and changed proportionally; ii) that the periaxonal space under the paranode was narrower than under the internode and did not exceed 3nm; and iii) that the periaxonal space is narrower at the paranode, but changes proportionally with the internodal periaxonal space (**Figure 6, S5**). While narrowing the periaxonal space at the paranode increased predicted conduction velocities compared to a theoretically uniform periaxonal space width, in each of these scenarios adjusting the node length and the internodal periaxonal space width to match the experimental values obtained following iTBS consistently slowed the simulated conduction speeds compared to the sham condition (**Figure 6, S5**), though the effect was not as strong when the paranodal periaxonal space width remained constant.

When we performed computation modelling to assess the impact that each ultrastructural adaptation measured could have on conduction velocity, we found that the iTBS-induced decrease in node length was predicted to slow conduction velocity, while the learning induced increase in node length was predicted to increase conduction velocity (**Figure 6; S5**). These data are consistent with the theoretical inverse U relationship between conduction speed and node length in cortical axons, when sodium channel density at the node of Ranvier remains constant (Arancibia-Carcamo *et al*., 2017). These earlier predictions indicate that at an average internode length of either ∼50μm (as per our CC data), or ∼37μm (as per our fimbria data), conduction velocity would be faster when node lengths were set to 1.5μm compared to 0.5μm, though further increasing node length to 3.5μm was predicted to markedly slow conduction speed when internode lengths were shorter than ∼50μm (Arancibia-Carcamo *et al*., 2017). In the context of iTBS, our computational modelling suggests that expanding the periaxonal space has a larger effect on action potential slowing than shortening the node length; however, following spatial learning, increasing node length and decreasing the width of the periaxonal space sped up action potential conduction to a similar extent. In both cases, altering node length and periaxonal space width had a synergistic effect, working together to slow or speed action potential conduction in the iTBS and learning simulations, respectively. These data are the first to demonstrate a synergistic relationship between node length and periaxonal space width in regulating the speed of action potential conduction along myelinated axons.

### Regulating conduction velocity is important for neural network function

It is likely that the dynamic modulation of action potential conduction velocity could be an adaptive mechanism to ensure the coincident arrival of action potentials at post-synaptic targets, and a mechanism that will allow circuits to adapt to the altered stimulation of specific circuits. Indeed, when we performed *ex vivo* CAP recordings in the CC of iTBS or sham stimulated mice we found that iTBS not only slowed CAP conduction along these axons, as our simulations predicted, but that the peak amplitude of the CAP response was increased and the half-width decreased, indicating a more synchronised arrival of action potentials at the recording electrode (**Figure 7**). At the network level, adjusting signal arrival at post-synaptic neurons could also affect the balance of excitation and inhibition, effectively altering their probability of firing, and may also determine whether potentiation or depression is induced (Feldman, 2012; Markram *et al*., 2012). The precise regulation of action potential conduction velocity and the firing frequency of neuronal populations is also important for synchronous or time-locked brain wave oscillations (Pajevic *et al*., 2014), which are associated with various cognitive functions including selective attention, information processing, sensory gating of information, learning, memory formation and consciousness (Ainsworth *et al*., 2012; Burgess *et al*., 2007; Buzsaki, 2006; Sirota *et al*., 2008). We propose that the combined plasticity of the periaxonal space and the nodes of Ranvier is a vital homeostatic mechanism for refining action potential conduction in response to altered neural circuit activity.

## Acknowledgments

We thank our colleagues at the University of Tasmania and Rowan Tweedale (Queensland Brain Institute, the University of Queensland) for constructive feedback on the manuscript and suggestions for improvement. We also thank Dr Lee Cossell and Prof David Atwell (University College London) for their advice on computational modelling, as well as Dr Carola Thoni and Mr Shane Rix (Lastek: Photonics Technology Solutions, Australia) for their assistance with STED imaging.

## Author contributions

CLC, KMY, RBJ, REP, KAP, JR and BE developed the project and wrote the manuscript. CLC, REP, KAP, MTC, LA, VO, RBJ and KMY carried out the experiments. KMY, JR, CLC, BE and RBJ obtained the funding. CLC, REP, MTC and RBJ performed the statistical analyses and generated the figures. KMY, ADT, GR, JR and CLC provided supervision.

## Methods

### Lead contact and materials availability

Further information and requests for resources and reagents should be directed to and will be fulfilled by the lead contact, Kaylene Young.

### Experimental model and subject details

All animal experiments were approved by the University of Tasmania Animal Ethics Committee and carried out in accordance with the Australian code of practice for the care and use of animals for scientific purposes. All wildtype and transgenic mice were maintained on a C57BL/6J background. Heterozygous *Plp-CreER* transgenic mice (Doerflinger *et al*., 2003) were crossed with heterozygous *Tau-lox-STOP-lox-mGFP-IRES-NLS-LacZ-pA* (*Tau-mGFP*) Cre-sensitive reporter mice (Hippenmeyer *et al*., 2005)to generate double heterozygous offspring for the fluorescent labeling and tracing of oligodendrocytes. Homozygous *Myrf* loxP-flanked exon 8 mice (*Myrf* ^*fl/fl*^) (Emery *et al*., 2009) were crossed with heterozygous *Pdgfra-CreER*^*TM*^ transgenic mice (Kang *et al*., 2010) or homozygous *Rosa26-YFP Cre-*sensitive reporter mice (Srinivas *et al*., 2001) to produce *Pdgfra-CreER*^*TM*^ *:: Myrf* ^*fl/fl*^ and *Myrf* ^*fl/fl*^ *:: Rosa26-YFP* offspring, respectively. These offspring were then intercrossed to generate *Pdgfra-CreER*^*TM*^ *:: Rosa26-YFP :: Myrf* ^*fl/fl*^ (Myrf ^fl/fl^) and *Rosa26-YFP :: Myrf* ^*fl/fl*^ (control) mice for experiments. Male and female mice were housed in same sex groups (2-4 per cage), in individually ventilated cages (Optimice®) on a 12 h light cycle (twilight phase starts 06:30, full lights on 07:00) at 21 ± 2°C with *ad libitum* access to standard rodent chow (Barrastoc rat and mouse pellets) and water. Experimental mice (P60-P90) weighed 18-35g at the start of experiments and were randomly assigned to each treatment, but care was taken to ensure littermates were represented across treatment groups.

### Method details

#### Transgenic lineage tracing and gene deletion

*Cre, Rosa26-YFP* and *Tau-mGFP* transgenes were detected by PCR as described by Cullen *et al*. (2019), and the *Myrf* ^*floxed*^ gene was detected as described by Emery *et al*. (2009). In brief, genomic DNA was extracted from ear biopsies by ethanol precipitation and PCR was performed using 50-100ng of gDNA with the following primer combinations: Cre 5’ CAGGT CTCAG GAGCT ATGTC CAATT TACTG ACCGTA and Cre 3’ GGTGT TATAAG CAATCC CCAGAA; GFP 5’ CCCTG AAGTTC ATCTG CACCAC and GFP 3’ TTCTC GTTGG GGTCT TTGCTC; Rosa26 wildtype 5’ AAAGT CGCTC TGAGT TGTTAT, Rosa26 wildtype 3’ GGAGC GGGAG AAATG GATATG and Rosa26 YFP 5’ GCGAA GAGTT TGTCC TCAACC; Myrf 5’ AGGAG TGTTG TGGGA AGTGG and Myrf 3’ CCCAG GCTGA AGATG GAATA.

To activate Cre-recombinase in oligodendrocyte precursor cells and induce targeted DNA recombination, Tamoxifen was dissolved in corn oil (40mg/ml) by sonication at 21°C for 2 h and administered to adult mice (P83) by oral gavage at a dose of 300mg tamoxifen/kg body weight daily for four consecutive days. *Plp-CreER :: Tau-mGFP* mice were given a single dose of 50mg/kg, 100mg/kg or 300mg/kg body weight to enable clear visualization of individual mGFP^+^ internodes (**Figure S1**).

#### Low intensity repetitive transcranial magnetic stimulation

Low intensity repetitive transcranial magnetic stimulation (Li-rTMS) was delivered as per Cullen *et al*. (2019). Briefly, 600 pulses of intermittent theta burst stimulation (iTBS; 192s) was delivered using a custom made 120mT circular coil designed for rodent stimulation (8mm outer diameter, iron core) (Tang *et al*., 2016a; Tang *et al*., 2016b). Stimulation parameters were controlled by a waveform generator (Agilent Technologies) connected to a bipolar voltage programmable power supply (KEPCO BOP 100-4M, TMG test equipment). Experiments were conducted at 100% maximum power output (100V) using custom monophasic waveforms (400μs rise time; Agilent Benchlink Waveform Builder). Mice were restrained using plastic body-contour shape restraint cones (0.5mm thick; Able Scientific). The coil was manually held over the midline of the head with the back of the coil positioned in line with the front of the ears (∼Bregma −3.0). Sham mice were positioned under the coil for 192s (as per iTBS), but no current was passed through the coil. Stimulation was carried out once daily, at the same time, for 14 consecutive days. Li-rTMS did not elicit observable behavioral changes in the mice during or immediately after stimulation.

#### Spatial learning

To induce spatial learning, adult (P60) male and female *Plp-CreER :: Tau-mGFP* or littermate control mice were administered a single dose of tamoxifen (50mg/kg), then handled daily for two weeks before being trained in an 8-arm radial arm maze (RAM) task (**Figure S3**) over 14 days. 5 days prior to RAM training, non-learning and learning mice had their access to normal mouse chow restricted to 6h per day but were given food rewards (Froot Loops® pieces) in their home cage. This food restriction protocol ensured that mice were maintained at ∼90% of their free feeding body weight and were motivated to seek out and consume the food rewards when available.

The RAM was carried out using the multi-maze system for mice (Ugo-Basile) in a radial 8-arm configuration with spatial cues placed on each of the surrounding walls, ∼30cm above the maze. RAM training consisted of two phases - a familiarization phase (days 1-3) and a learning phase (days 4-14) (**Figure S3**), and each mouse underwent 3 trials per day, with 60 minutes between each trial. During the familiarization phase, all arms of the RAM were closed off and an individual mouse was placed in the octagonal center of the maze with a single Froot Loop® (cut into 8 approximately equal sized pieces) for 10 minutes. No-learning control mice were exposed to the familiarization phase for the 14 days of the task, to ensure that they were subjected to the same environment and an equivalent level of handling and that they also received Froot Loops®, but did not learn the RAM task (**Figure S3A**). Learning mice proceeded to undertake a learning phase, in which the 8 pieces of Froot Loop® were distributed, so that one piece was placed at the end of each arm of the RAM. An individual mouse was placed in the center of the maze and could explore the maze for 10 minutes (**Figure S3B**). Over the next 11 days, the mice learned that each arm contained a single food reward and that repeat entries would not result in another reward. Therefore, repeated entries into an arm in which the food reward had already been consumed was counted as an error, and the average number of errors made per trial was quantified as a measure of learning (**Figure S3C**). 24 hours after the final trial, mice were perfusion fixed and their brains collected and prepared for either immunohistochemistry or transmission electron microscopy.

#### Tissue preparation and immunohistochemistry

Mice were perfusion-fixed with 4% paraformaldehyde (PFA; Sigma) (w/v) in phosphate buffered saline (PBS). Brains were cut into 2mm-thick coronal slices using a 1mm brain matrix (Kent Scientific) before being post-fixed in 4% PFA at 21°C for 90 minutes. Tissue was cryoprotected overnight in 20% (w/v) sucrose (Sigma) in PBS and snap frozen in OCT (ThermoFisher) for storage at −80°C. 30μm coronal brain cryosections, containing the primary motor cortex and underlying corpus callosum (∼Bregma +0.5) or the dorsal region of the fimbria (∼Bregma −1.5), were collected and processed as floating sections (Cullen *et al*., 2019). Primary and secondary antibodies were diluted in PBS blocking solution [0.1% (v/v) Triton X-100 and 10% fetal calf serum in PBS] and applied to sections overnight at 4°C, unless staining involved the use of mouse anti-CC1 (1:100 Calbiochem), in which case antibodies were diluted in Tris buffered saline (TBS) blocking solution [0.1% (v/v) Triton X-100 and 10% fetal calf serum in TBS]. Primary antibodies included goat anti-PDGFRα (1:200; R&D Systems), rabbit anti-OLIG2 (1:400 Millipore), rat anti-GFP (1:2000; Nacalai Tesque), rabbit anti-NaV_1.6_ (1:500 Alomone Labs), mouse anti-CASPR (Clone K65/35; 1:200 NeuroMab). Secondary antibodies, which were conjugated to AlexaFluor −488, −568 or −647 (Invitrogen) were donkey anti-goat (1:1000), donkey anti-rabbit (1:1000), donkey anti-mouse (1:1000), and donkey anti-rat (1:500). Nuclei were labeled using Hoechst 33342 (1:1000; Invitrogen).

#### Confocal microscopy and image quantification

Confocal images were collected using an UltraView Nikon Ti Microscope with Volocity Software (Perkin Elmer). High magnification images (z-spacing of 0.5-2μm) were collected using standard excitation and emission filters for DAPI, FITC (AlexaFluor-488), TRITC (AlexaFluor-568) and CY5 (AlexaFluor-647), then stitched together to make a composite image of a defined region of interest. To quantify internode number and length for oligodendrocytes within the primary motor cortex (M1), high magnification images (40x objective) were collected through individual mGFP-labeled cortical oligodendrocytes (0.5μm z-steps) that had a visible cell body. To quantify internodes in the corpus callosum (CC) and hippocampal fimbria, high magnification images (60x objective) were collected (0.5μm z-steps) and used to identify individual mGFP-labeled internodes that were flanked by CASPR^+^ paranodes. To measure node of Ranvier (Na_v_1.6) and paranode (CASPR) length, high magnification (100x) single z-plane confocal images were collected from M1, the CC and the fimbria. Node and paranode lengths were only measured when a node and its flanking paranodes were intact within the single z-plane. For quantification of cell number, low magnification (20x objective) confocal z-stacks (2 μm spacing) were collected through M1 and the CC and stitched together to make a composite image of a defined region of interest. All image analysis was carried out using Image J (NIH) by a researcher blind to experimental treatment.

#### Stimulated emission depletion (STED) microscopy

30μm coronal cryosections containing the corpus callosum (∼Bregma +0.5) were collected and prepared as floating sections. Rabbit anti-Na_v_1.6 (1:500 Alomone Labs), mouse anti-CASPR (Clone K65/35; 1:200 NeuroMab) primary antibodies were diluted in PBS blocking solution [0.1% (v/v) Triton X-100 and 10% fetal calf serum in PBS] and applied to sections overnight at 4°C. The sections were washed in PBS (3 x 10min) before overnight application (4°C) of goat anti-mouse STAR Red (1:500, Abberior) and goat anti-rabbit STAR Orange (1:500, Abberior) secondary antibodies. The sections were mounted in antifade liquid mounting media (Abberior) and covered in a 170 μm thick glass coverslip (ProSciTech, cat # EMS72291-06).

STED imaging was performed using a two-color Abberior STEDYCON system (Abberior Instruments GmbH) attached to a Nikon NiE confocal microscope equipped with 405nm, 488 nm, 561 nm, and 640 nm pulsed excitation lasers, a pulsed 775 nm STED laser and a 100x oil immersion objective lens (N.A 1.4). Images were acquired using Abberior STEDYCON smart control software. For all images the pixel size and dwell time were kept consistent at 20nm and 10μs, respectively. 561nm and 640nm excitation lasers were set to 10% power but STED laser power was optimally set to 100% (STAR orange, 561nm) or 56.2% (STAR red, 640nm). Single z-plane STED images were collected from the CC to enable the precise, high-resolution visualization of nodes of Ranvier (Na_v_1.6) and their abutting paranodes (CASPR).

#### Transmission electron microscopy

Following 14 days of iTBS, sham stimulation or RAM training mice (P105 or P89) were perfused with Karnovsky’s fixative (2.5% glutaraldehyde, 2% PFA, 0.25mM CaCl_2_, 0.5mM MgCl_2_ in 0.1M sodium cacodylate buffer). Brains were cut into 2mm-thick coronal slices using a 1mm brain matrix (Kent Scientific) and post-fixed in Karnovsky’s fixative for 2h at 21°C. The tissue blocks were rinsed and stored in 0.1M sodium cacodylate buffer overnight. The CC or hippocampal fimbria were dissected and incubated in 1% osmium tetroxide / 1.5% potassium ferricyanide [OsO_4_ / K_3_Fe(III)(CN)_6_] in 0.1M sodium cacodylate buffer in the dark for 2h at 4°C, before being dehydrated in ethanol and propylene oxide, and embedded in Epon812 resin. Ultrathin 70nm sections were cut using a Leica Ultra-cut UCT7 and stained with uranyl acetate and lead citrate. High resolution electron microscopy imaging was done at 80kV on a JEOL 1400-Flash (CC) or a Hitachi HT7700 (fimbria) transmission electron microscope. Sectioning, imaging and image analysis was carried out by an experimenter blind to the treatment group.

Image analysis was carried out using Image J (NIH). The proportion of myelinated axons and the g-ratio of myelinated axons [axon diameter / (axon + myelin diameter)] were measured from at least 100 axons from 5 images per animal. The number of myelin wraps was quantified by counting major dense lines for a minimum of 15 transected axons per mouse, from n=3 mice per treatment group. The average thickness of myelin wraps per axon, the width of the adaxonal inner tongue membrane and the width of the periaxonal space were measured from axons ensheathed by myelin that was a minimum of five wraps thick (≥10 transected axons per mouse, from n=3 mice per treatment group).

#### Mathematical modeling

In order to evaluate the effect on action potential propagation of experimentally observed microscopic changes in node length and myelin structure [data derived from the population mean from n=3 animals, (**Figures 1, 2, 5 and S2**) for iTBS experiments; and n=3-4 animals, (**Figures 3, 5 and S3**) for RAM experiments], we further adapted the mathematical model of action potential propagation in myelinated axons proposed by Richardson and colleagues [model ‘C’, their Fig. 1; (Bakiri *et al*., 2011; Richardson *et al*., 2000)]. A recent MATLAB (The MathWorks) implementation of that model by Cossell and colleagues can be downloaded from GitHub (https://github.com/AttwellLab/MyelinatedAxonModel) (Arancibia-Carcamo *et al*., 2017; Bakiri *et al*., 2011; Ford *et al*., 2015; Young *et al*., 2013). That package was downloaded in June 2018 and run on MATLAB R2016b.

The mathematical description of ion channels at nodes of Ranvier follows the Hodgkin-Huxley formalism. Briefly, nodes express three types of ion channels, a fast sodium channel 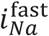 responsible for the initiation of action potentials, a persistent sodium channel 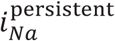, and a slow potassium channel 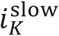 responsible for the termination of action potentials. The kinetics of the three currents is derived from McIntyre *et al*. (2002). Briefly, 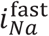 is written:

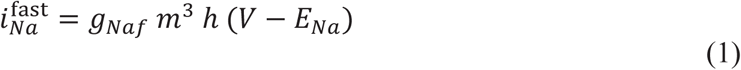

with *g*_*Naf*_ the current conductance, *V* the membrane voltage at the node, *E*_*Na*_ = 60mv the reversal potential for sodium ions, and *m* and *h* some gating variables. Following the Hodgkin-Huxley formalism, each gating variable *x* in the model follows the generic equation:

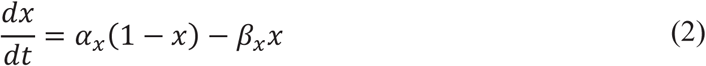

with *α* and *β* some functions of *V*. For 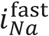, *α* and *β* are given by:

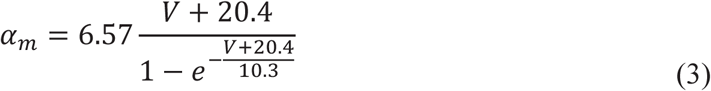

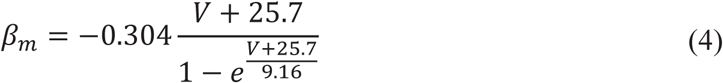

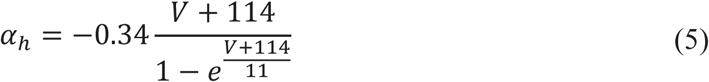

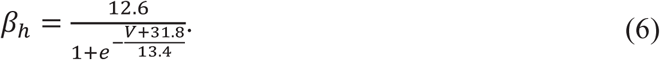

The persistent sodium current 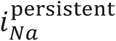 is given by:

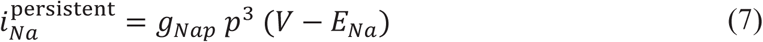

with:

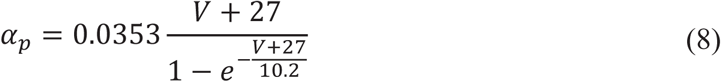

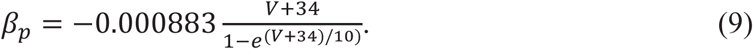

The slow potassium current 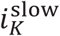 is given by:

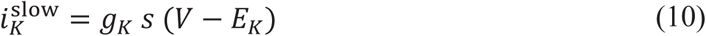

with *E*_*k*_ = −90mv the reversal potential for potassium ions and:

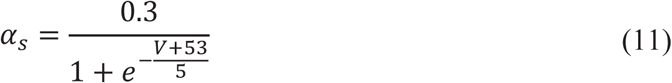

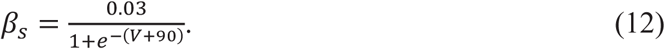

Finally, all membranes contain a leak current given by:

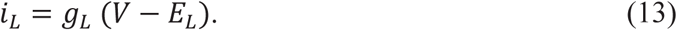

The Q10 for gates *m, h, p* and *s* are 2.2, 2.9, 2.2 and 3.0 respectively, described at 36°C [note that this has the effect of slowing down the kinetics of gates *m, h* and *p* with respect to McIntyre *et al*. (2002)]. All the other parameters of the model are given in **Table S1** below. The same framework was used to simulate both *corpus callosum* and *fimbria* axons but with different parameter sets (see **Table S1**). Numerical values were chosen to match those observed in experiments or adapted from (Arancibia-Carcamo *et al*., 2017)). Myelin thickness was automatically calculated using:

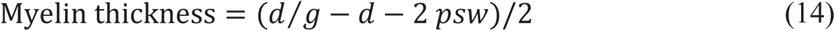

with *d* the axon diameter, *g* the g-ratio and *psw* the periaxonal space width. Myelin lamella periodicity was taken as myelin thickness divided by 6.5 so that the number of wraps is 7 in all conditions as observed experimentally (see **Figure 5**), assuming that the extracellular space between myelin lamellae comprises part of the periodicity and to account for the fact that there is no extracellular space contributing to the total width of the myelin on the most external lamella.

Unless stated otherwise, simulations were run using a time step of 0.1μs and 51 nodes. Internode segments were chosen to be <1μm (0.98μm; N = 52 segments per internode) and we verified that this was sufficient to reach convergence for the conduction velocity over the whole range of simulated axons (**Figure S6**). Action potentials were triggered by a square pulse of 0.5nA lasting 10μs. Ion channels at juxtaparanodes were not modelled, as is common in the field. When altering the length of the node of Ranvier (see below), the density of ion channels at the node was taken to be constant [see (Arancibia-Carcamo *et al*., 2017) for a systematic discussion of how this affects action potential conduction velocity].

To evaluate individually the effect of a node length reduction or the myelin sheath alterations on conduction velocity, we initially ran four sets of simulations. First, we used a parameter set matching the observations obtained in the sham condition (column ‘Sham’ in **Table S1**). We then ran the same simulations after reducing node lengths (‘Short nodes’). Third, we ran simulations modifying the myelin sheath but keeping node length as per the sham condition (‘Alt. myelin’). Finally, we ran a simulation with a fourth set of parameters implementing all the experimental changes observed following iTBS, i.e. a reduction in node length and alterations in the myelin sheath (‘iTBS’). Simulations were run at 21°C and at 37°C (**Figure 6, S5**). We additionally investigated three different scenarios for conduction at the paranode. The periaxonal space width at the paranode was taken to be either [i] equal to the periaxonal space width in the internode; [ii] equal to the periaxonal space width under the internode if that is less than 3nm, but to be at most 3nm otherwise; or [iii] equal to half the periaxonal space width in the internode (**Figure 6, S5**). Unless otherwise specified, scenario [i] is in use. Each paranode was taken to be 2 segments long (1.96μm long).

To evaluate the functional consequence of myelin alterations, we additionally ran a set of simulations varying the periaxonal space width from 0 to 20nm at both 21°C and 37°C (**Figure 6**). These simulations show that at 37°C, the periaxonal space can tune action potential conduction velocity between 4.36m/s (psw = 0nm) and 1.25m/s (psw = 20nm; **Figure 6**). These numbers illustrate how potent and elegant this mechanism is, as it can speed up or slow down action potential conduction by a factor of 3.5 by making minor adjustments to the structure of myelinated axons. The functional consequences of this change to propagation speed at 37°C is to alter the arrival time of action potentials by 6ms over a distance of 1cm (**Figure 6**), enough to alter learning via spike-timing dependent plasticity for instance.

#### Compound action potential recordings

Compound action potential (CAP) recording and conduction-velocity measurement procedures were adapted from (Crawford *et al*., 2009). Briefly, the day after stimulation was complete, sham or iTBS-treated mice were killed by cervical dislocation and their brains rapidly dissected into ice-cold sucrose solution containing: 75 mM sucrose, 87 mM NaCl, 2.5 mM KCl, 1.25 mM NaH_2_PO_4_, 25 mM NaHCO_3_, 7 mM MgCl_2_, and 0.95 mM CaCl_2_. Coronal vibratome sections (400 μm; 2-3 per animal spanning Bregma +0.8 and −0.2) were generated using a Leica VT1200s vibratome and incubated at ∼32°C for 45 min in artificial cerebral spinal fluid (ACSF) containing 119 mM NaCl, 1.6 mM KCl, 1 mM NaH_2_PO4, 26.2 mM NaHCO_3_, 1.4 mM MgCl_2_, 2.4 mM CaCl_2_ and 11 mM glucose (300 ± 5 mOsm / kg), before being transferred to ∼21°C ACSF saturated with 95% O_2_ / 5% CO_2_.

CAPs were evoked by constant current, stimulus-isolated, square wave pulses (200 ms duration, delivered at 0.2 Hz), using a tungsten bipolar matrix stimulating electrode (FHC; MX21AEW), and detected using glass recording electrodes (1-3 MΩ) filled with 3M NaCl. To quantify CAP amplitude, the asymptotic maximum for the short-latency negative peak (myelinated peak, M; **Figure 7**) was first determined by placing the stimulating and recording electrodes 1mm apart and varying the intensity of stimulus pulses (0.3–4.0 mA) using an external stimulus isolator (ISO-STIM 1D) before recording at 80% maximum stimulation. To enhance the signal-to-noise ratio, all quantitative electrophysiological analyses were conducted on waveforms that were the average of eight successive sweeps, amplified and filtered (10 kHz low pass bessel) using an Axopatch 200B amplifier (Molecular Devices), digitized at 100 kHz and stored on disk for offline analysis.

The conduction velocity of myelinated (M) and unmyelinated (UM) axons in the CC was estimated by changing the distance between the stimulating and recording electrodes from 1 to 3 mm, while holding the stimulus intensity constant (80% maximum). The peak latency of the M and UM axons was measured at each point and graphed relative to the distance separating the electrodes. A linear regression analysis was then performed to yield a slope that is the inverse of the velocity for each brain slice. The average velocity for both CAP components (M, UM) was then determined for each animal (n=7 per group) and this value was used for statistical comparison.

### Statistical analyses

The number of mice analyzed in each group (*n*) or the number of cells, axons, nodes or internodes is indicated in the corresponding figure legends. Data distributions are presented as cumulative distribution plots and as violin plots with the median and interquartile range indicated. Data averaged per animal are presented as mean ± SD with all data points shown. All statistical analyses were performed using Prism 8 (GraphPad Software). Data comparing two groups at a single time point were analyzed using a parametric two-tailed t-test (*n* = mouse) or a non-parametric Mann-Whitney U test (MWU; *n* = node, paranode, internode or axon). Cumulative distribution data were analyzed using a Kolmogorov–Smirnov (KS) test. Cell counts for lineage tracing, and compound action potential data were analyzed using a 2-way ANOVA with Bonferroni post-test. Learning in the RAM was analyzed using a repeated measures one-way ANOVA with a Geisser-Greenhouse correction to ensure equal variability and sphericity was not assumed, followed by a Bonferroni post-test. ANOVA main effects are given in the corresponding figure legends.

### Data code and availability

Data for computational modelling are available from GitHub (https://github.com/JolivetLab).

### Key resources table

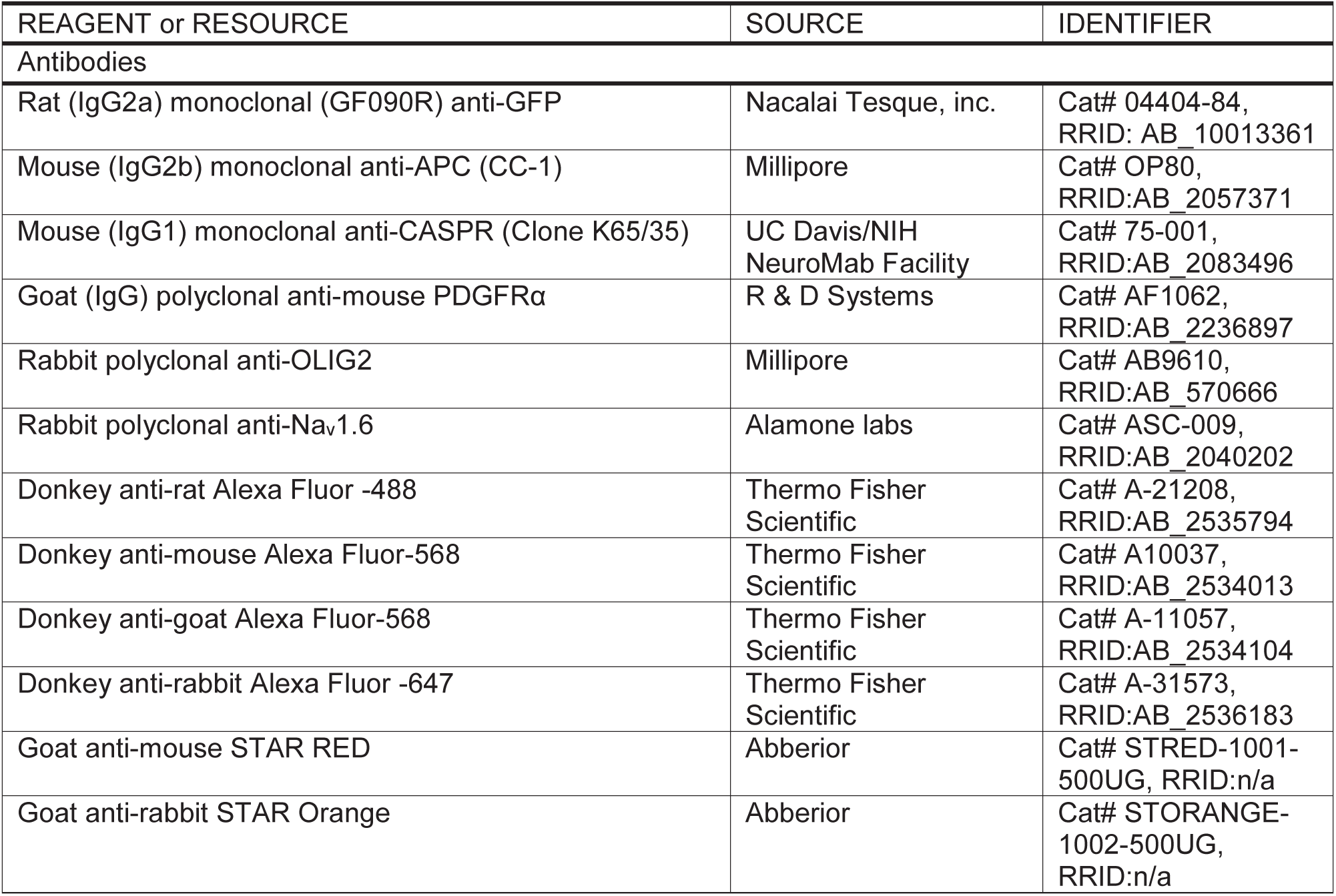
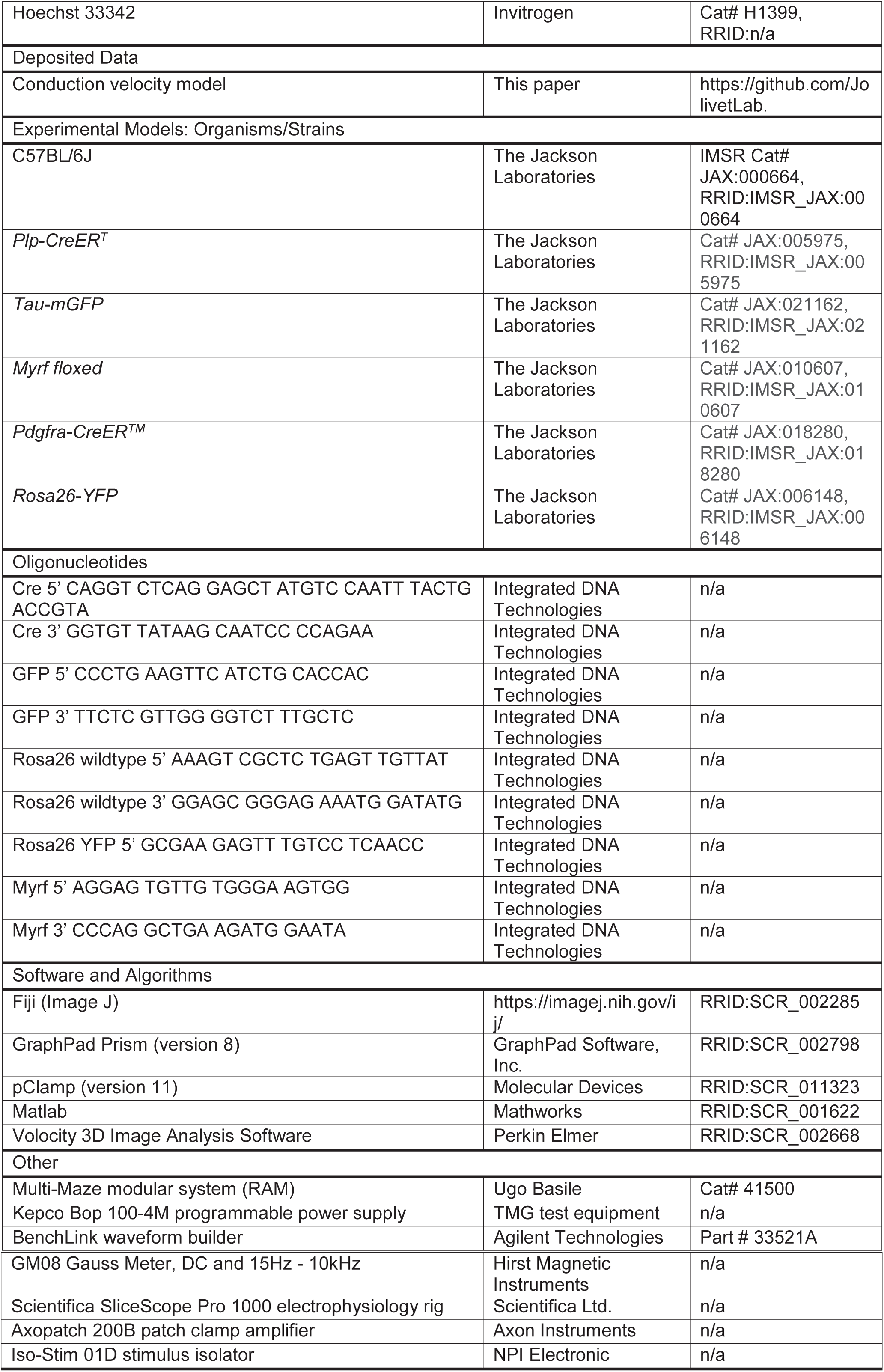

## Supplementary material

**Figure S1.**
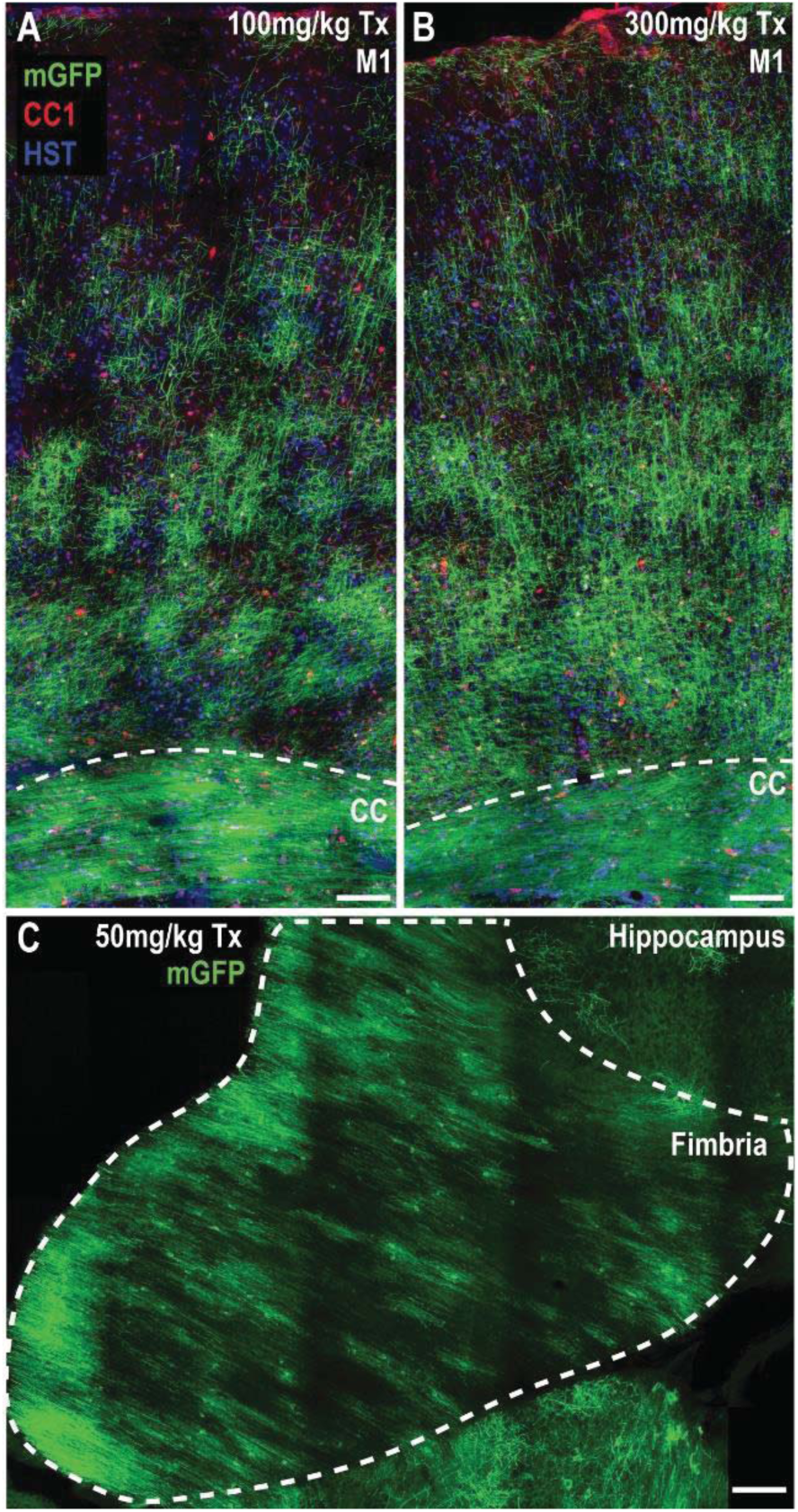
Mature, myelinating oligodendrocytes become mGFP-labeled following Tamoxifen administration to *PlpCreER :: Tau-mGFP* transgenic mice **A**) Confocal image of M1 and the CC of a P80+15 *Plp-CreER :: Tau-mGFP* mouse that received a single 100mg/kg dose of Tamoxifen (Tx), with its brain subsequently being stained to detect GFP (green), the oligodendrocyte marker CC1 (red) and Hoescht 33342 (blue). **B**) Confocal image of M1 and the CC of a P80+15 *Plp-CreER :: Tau-mGFP* mouse that received a single 300mg/kg dose of Tamoxifen, with its brain being stained to detect GFP (green), CC1 (red) and Hoescht 33342 (blue). In mice that received only 100mg/kg of Tamoxifen, it was possible to discern individual GFP-labeled CC1^+^ oligodendrocytes and their associated myelin internodes in the cortical layers. It was not possible to distinguish individual GFP-labeled CC1^+^ oligodendrocytes and their associated internodes within the CC of *PlpCreER ::Tau-mGFP* mice that received 100 or 300 mg/kg Tamoxifen. **C**) Confocal image of the hippocampal fimbria of a P60+15 *Plp-CreER :: Tau-mGFP* mouse that received a single 50mg/kg dose of Tx. It was not possible to reliably attribute mGFP^+^ internodes to a single oligodendrocyte within the fimbria. Scale bars represent 100μm (A-B), 75μm (C).

**Figure S2.**
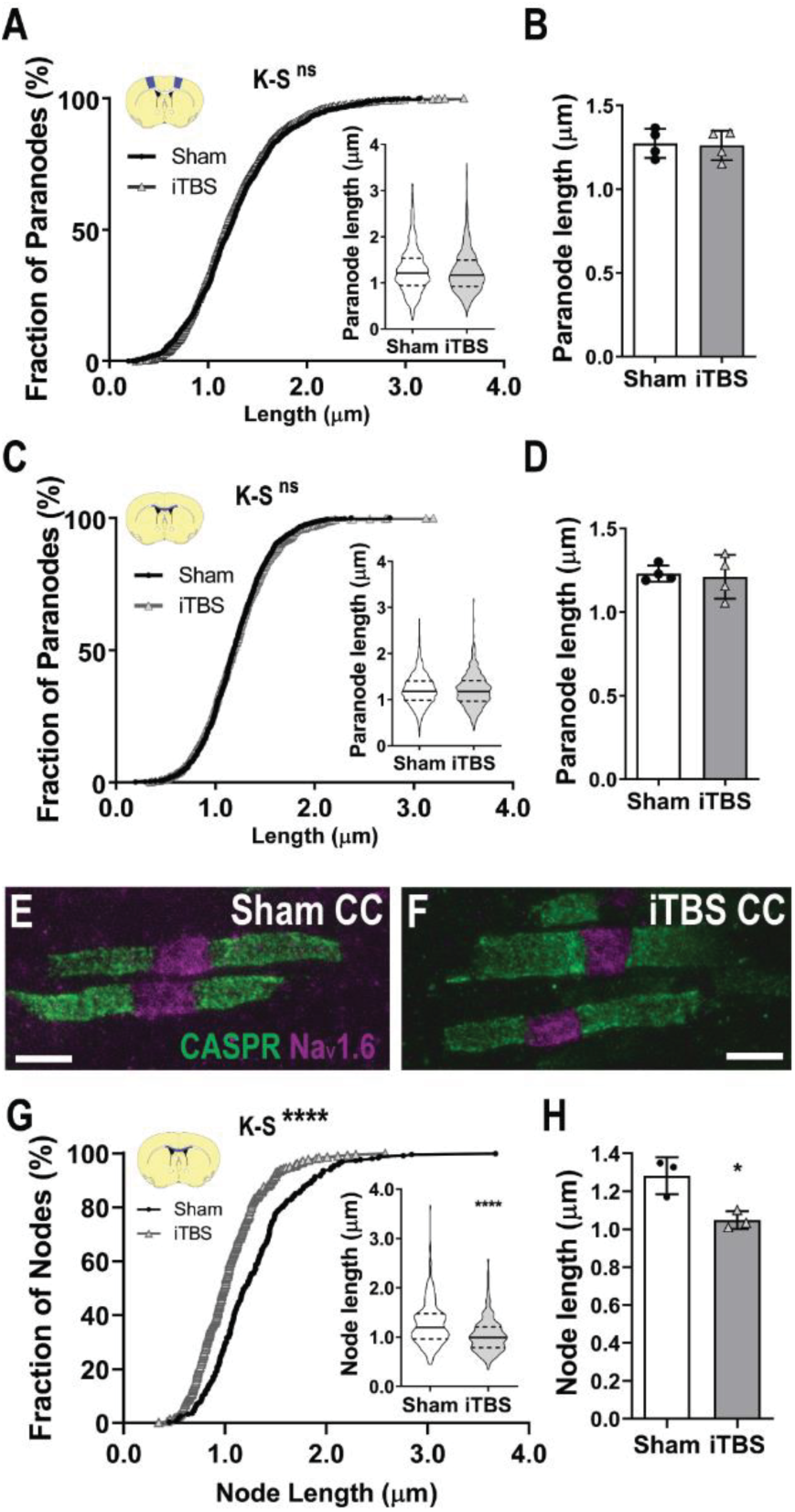
iTBS does not alter paranode length. **A**) Cumulative paranode length distribution in M1 [1268 sham (black circles) and 1335 iTBS (gray triangles) paranodes; K-S test, K-S D= 0.053, p=0.054; Inset, violin plot of paranode length, MWU test p=0.06]. **B**) Average M1 paranode length per individual sham (white) and iTBS (gray) treated mouse [n=4 mice per group, t-test, t=0.20, p=0.84]. **C**) Cumulative paranode length distribution in the CC [1099 sham and 1181 iTBS paranodes; K-S test, K-S D=0.03, p=0.65; Inset, violin plot of paranode length, MWU test p=0.73] of mice receiving sham-stimulation and iTBS. **D**) Average CC paranode length per sham and iTBS stimulated animal [n=4 per group, t-test, t=0.28, p=0.78]. **E-F**) Representative STED image of nodes of Ranvier (Na_v_1.6; magenta) and paranodes (CASPR; green) from the CC of sham (**E**) and iTBS (**F**) mice. **G**) Cumulative distribution plot of node length in the STED imaged CC of sham and iTBS treated mice [269 sham and 325 iTBS nodes; K-S test, K-S D=0.27, p<0.0001; Inset, violin plot of paranode length, MWU test p<0.0001]. **H**) Average CC node length (STED imaged) per sham and iTBS treated animal [n=3 per group, t-test, t=3.73, p=0.02]. Violin plots show the median (solid line) and interquartile range (dashed lines). Bars show mean ± SD. Scale bars = 1μm.

**Figure S3.**
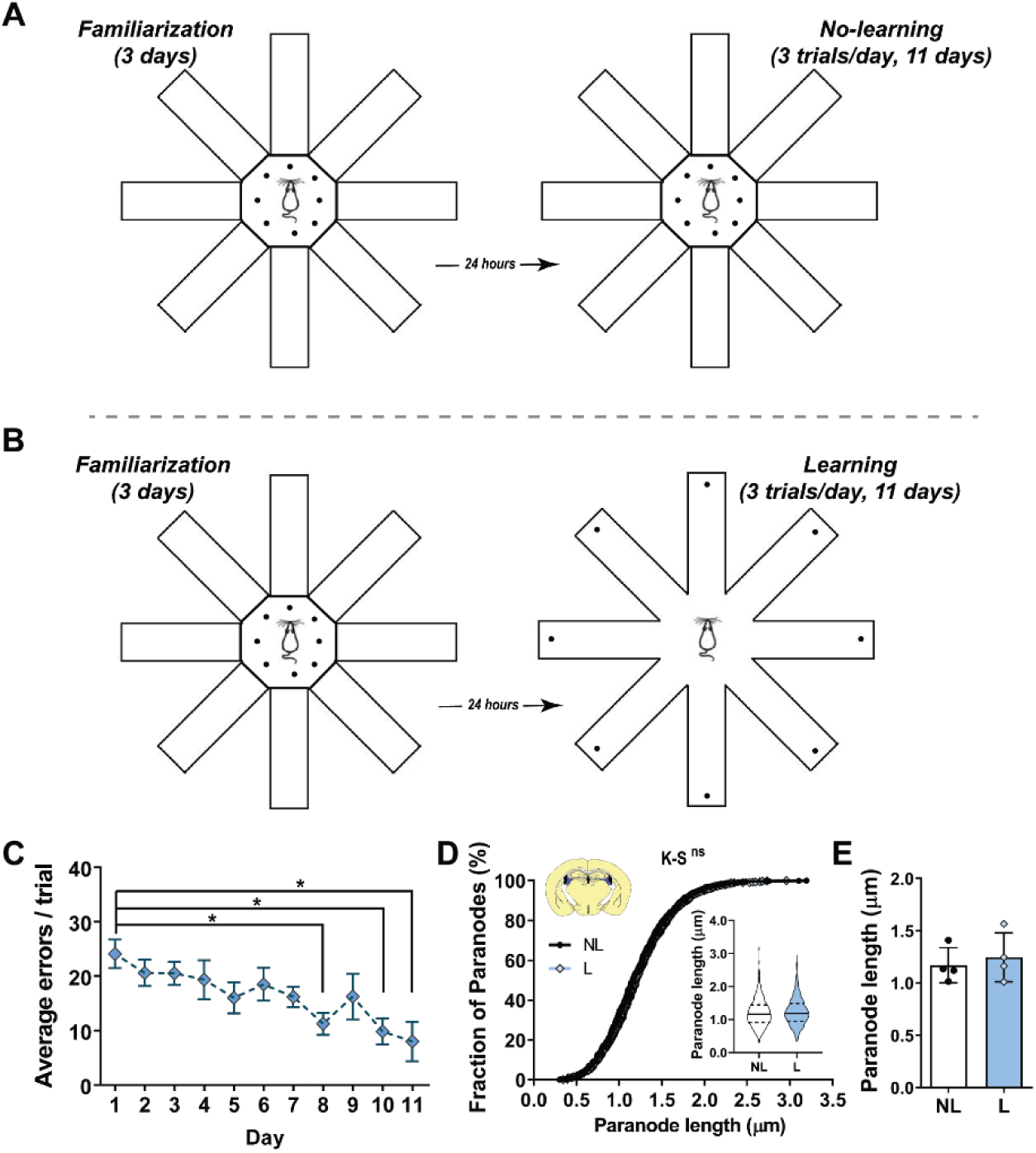
Spatial learning does not change paranode length in the fimbria. **A**) Schematic outlining the radial arm maze (RAM) procedure during the first 3 days of familiarization to the maze and over the following 11 days for no-learning control mice. Each dot represents a piece of Froot Loop® that was used as a food reward stimulus and placed in the center of the maze for each trial over a total of 14 days. **B**) Schematic outlining the procedure during the familarization phase (first 3 days) and learning phase (last 11 days) of the RAM spatial learning task. Each dot represents a piece of Froot Loop® that was used as a food reward stimulus. Food rewards were placed in the center of the maze during the familiarization phase, but a single piece of Froot Loop® was placed at the end of each arm during each trial of the learning phase. Mice learned to enter each arm and consume the single food reward only once during a trial, with repeated entries into an arm in which the food reward was already consumed being recorded as a recall error. **C**) Quantification of the average number of errors made across 3 trials each day by mice learning the RAM task. Over the course of 11 days the mice progressively learned the task and made fewer errors [RM one-way ANOVA with Bonferroni post-test: F(1.84, 5.521) = 5.62, p=0.048. Mean ± SEM for n=4 mice]. **D**) Cumulative paranode length distribution in the fimbria of mice that underwent no-learning (NL; black dots) or learning (L; blue diamonds) in the RAM [682 no-learning and 724 learning paranodes; K-S test, K-S D= 0.081, p=0.019; Inset, violin plot of paranode length, MWU test p=0.007]. **E**) Average fimbria paranode length in NL (white) and L (blue) mice [n=4 mice per group, t-test, t=0.52, p=0.61]. Violin plots show the median (solid line) and interquartile range (dashed lines). Bars show mean ± SD. *p<0.05

**Figure S4.**
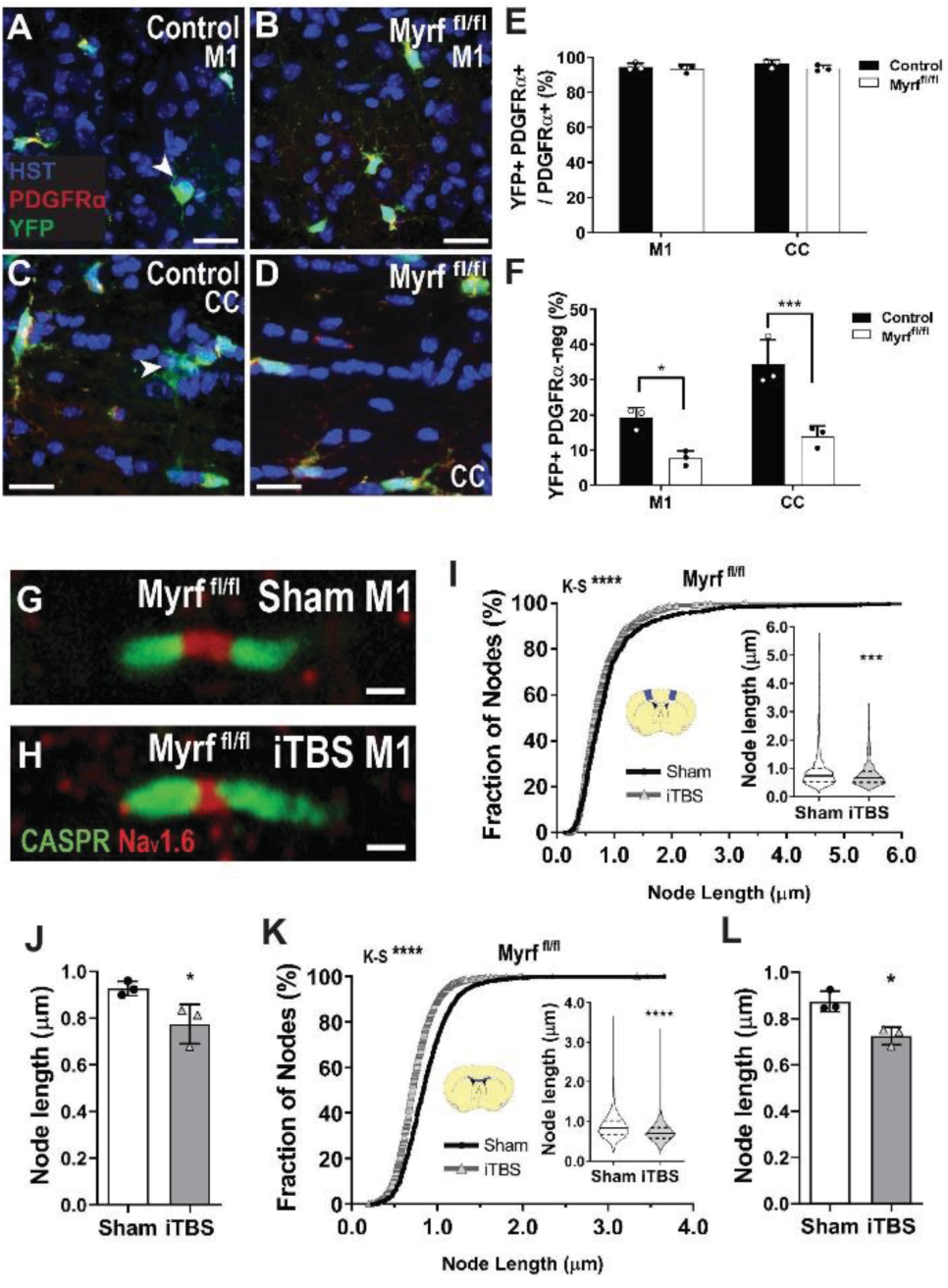
Preventing oligodendrogenesis does not affect iTBS induced node shortening. Tamoxifen was administered to P60 control (*Pdgfrα-CreER*^*TM*^*:: Rosa26-YFP*) and Myrf^fl/fl^ (*Pdgfrα-CreER*^*TM*^ *:: Rosa26-YFP :: Myrf* ^*fl/fl*^) mice to turn on expression of YFP in oligodendrocyte precursor cells (OPCs) and allow the lineage tracing of their progeny. Tamoxifen also resulted in the conditional deletion of *Myrf* from adult OPCs. **A-D**) Compressed confocal image from M1 (**A-B**) and CC (**C-D**) of P60+30 control (**A, C**) and Myrf^fl/fl^ (**B, D**) mice stained to detect PDGFRα (red), YFP (green) and Hoescht 33342 (blue). **E**) The proportion of PDGFRα+ OPCs that underwent recombination and become YFP-labeled in M1 and the CC of P60+30 control (black bars, white circles) and Myrf-deleted (white bars, black circles) mice [2-way ANOVA: region F(1,8)=0.57, p=0.46, gene F(1,8)=2.49, p=0.15, interaction F(1,8)=0.33, p=0.57. Mean ± SD. n=3 mice per group]. **F**) Quantification of the proportion of PDGFRα-neg YFP-labeled (and OLIG2^+^) newly differentiated oligodendrocytes in M1 and the CC [2-way ANOVA with Bonferroni post-test: region F(1,8)=19.86, p=0.0021, gene F(1,8)=43.56, p=0.0002, interaction F(1,8)=3.56, p=0.095]. **G-H**) Node of Ranvier (Na_v_1.6; red) in M1 of *Myrf*-deleted mice (*Pdgfrα-CreER*^*TM*^*::Myrf* ^*fl/fl*^) after 14 days of sham-stimulation (**G**) or iTBS (**H**). **I-J**) Cumulative node length distribution in M1 [**C**; 620 sham (black circles) and 727 iTBS (gray triangles) nodes; Kolmogorov-Smirnov (K-S) D=0.12, p<0.0001; inset violin plot of node length, Mann Whitney U (MWU) test, p<0.0001] and average node length per sham (white) or iTBS (gray) treated mice (**J**); [n=3 mice per group, t-test, t=2.93, p=0.04]. **K-L**) Cumulative node length distribution in the CC (**K**) [1903 sham and 1670 iTBS nodes; K-S D=0.17, p<0.0001; inset violin plot of node length, MWU test, p<0.0001] and the average node length per animal (**L**) of *Myrf*-deleted mice after sham-stimulation or iTBS [n=3 mice per group, t-test, t=4.453, p=0.01]. Arrows indicate newborn oligodendrocytes. Violin plots denote the median (solid line) and interquartile range (dashed lines). *p<0.05, ***p<0.001, ****p<0.0001. Scale bars represent 13μm (A-D), 1μm (G-H).

**Figure S5.**
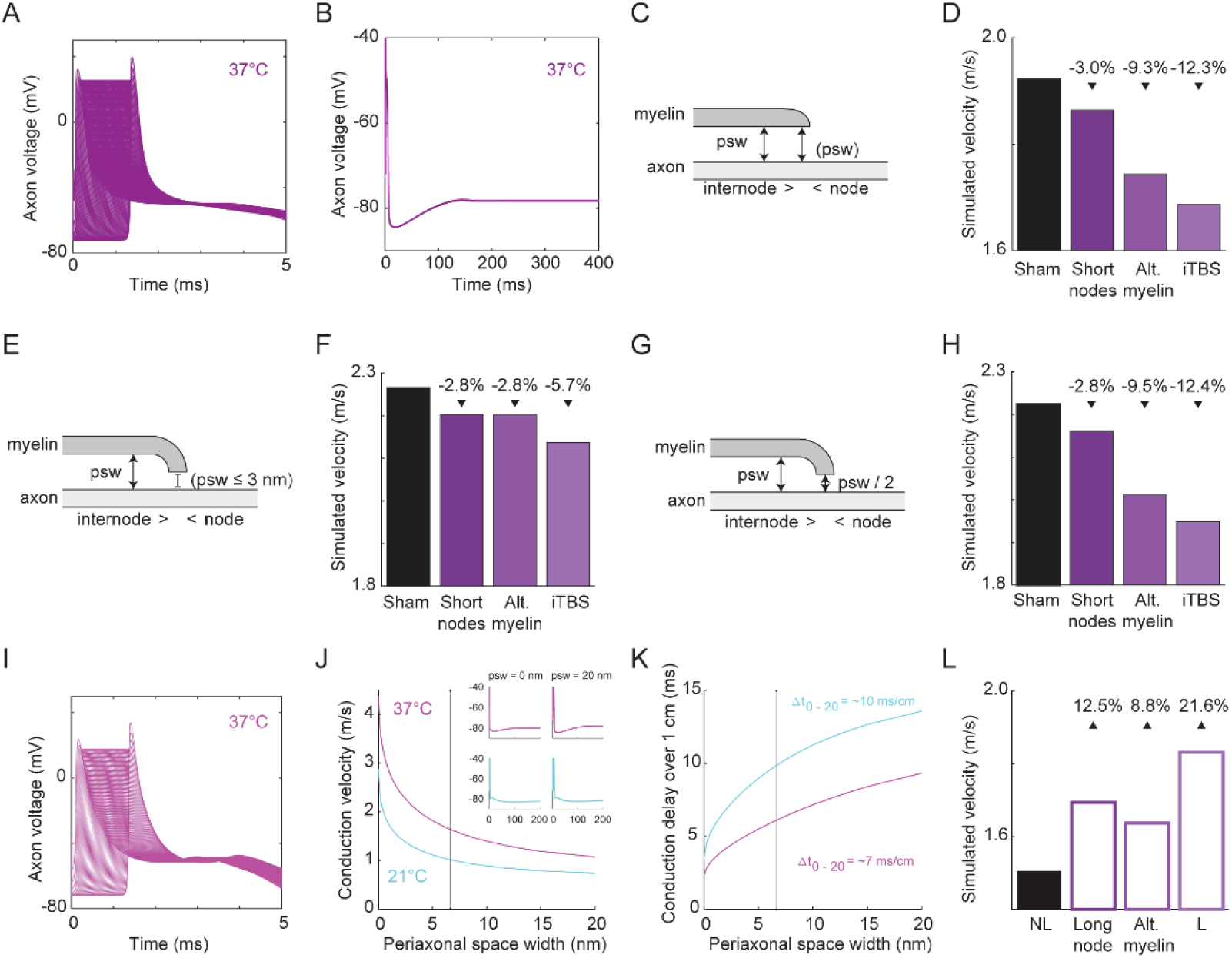
Mathematical simulation of conduction in myelinated central nervous system axons **A**) Action potentials simulated at consecutive nodes in the corpus callosum at 37°C. **B**) The extended time-course of action potentials generated by the model at 37°C. **C**) Schematic showing model parameters in which periaxonal space width (*psw*) at the paranode is uniform to internode *psw*. **D**) Predicted conduction velocity of a sham stimulated (white) axon versus an axon with either the node length shortened, the periaxonal space widened or both (iTBS, shades of magenta) using the model depicted in (C). **E**) Schematic showing model parameters in which *psw* at the paranode is set to ≤ 3nm. **F**) Predicted conduction velocity of a sham stimulated axon versus an axon with either the node length shortened, periaxonal space widened or both (iTBS) using the model depicted in (E). **G**) Schematic showing model parameters in which *psw* at the paranode is set to half the width in the internode. **H**) Predicted conduction velocity of a sham stimulated axon versus an axon with either the node length shortened, periaxonal space widened or both (iTBS) using the model depicted in (G). **I**) Action potentials simulated at consecutive nodes in the fimbria at 37°C. **J**) Simulated conduction velocity of fimbria axons relative to periaxonal space width (*psw*) at 37°C (light magenta) and 21°C (cyan), solid line indicates average *psw* following iTBS, insets show action potential waveforms at the extremities of the tested range (*psw* = 0 or 20nm) at 21°C and 37°C. **K**) Conduction delay over 1cm relative to *psw* in the fimbria at 21°C and 37°C. **L**) Predicted conduction velocity of a no-learning (NL) control axon within the fimbria at 37°C (white) versus an axon with either longer nodes, a narrower periaxonal space or both (learning, L).

**Figure S6.**
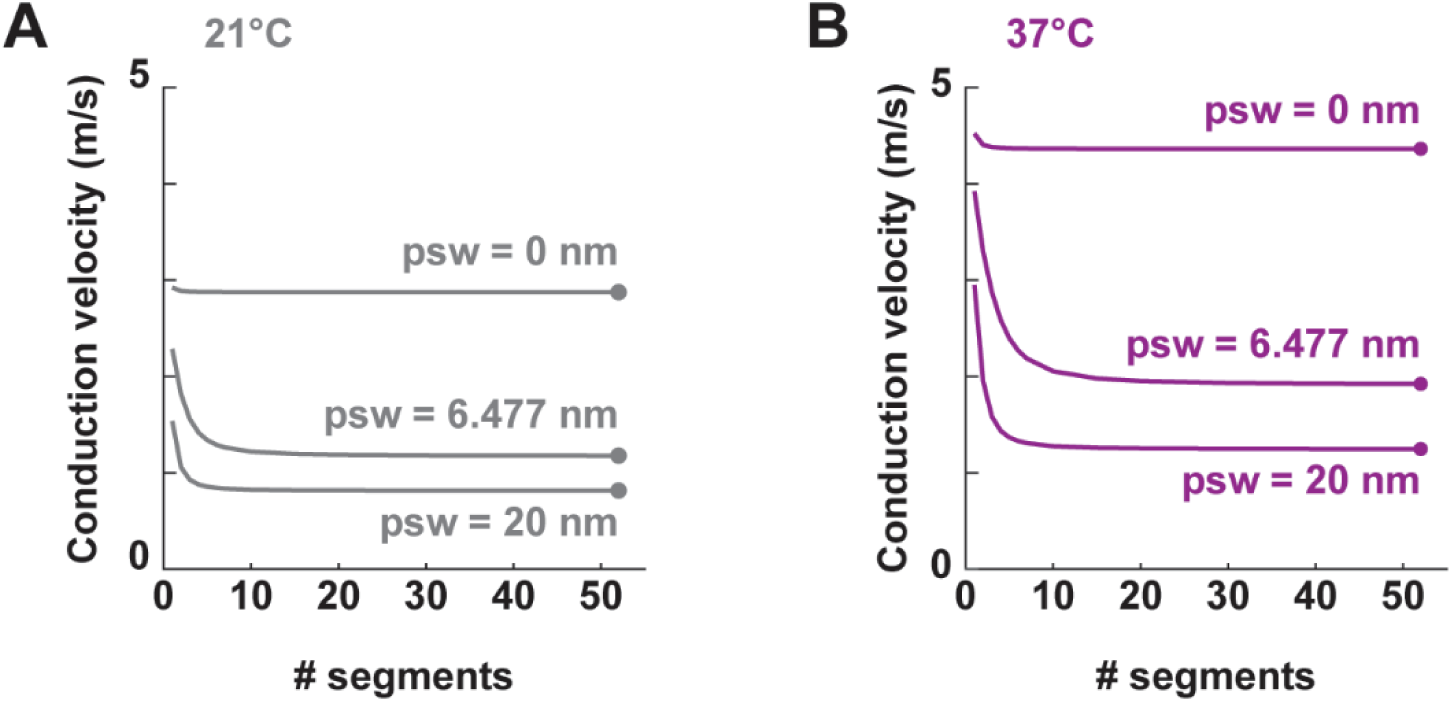
Controls for the convergence of conduction velocity in mathematical simulations Conduction velocity simulations were run at 21°C (**A**) or 37°C (**B**), inputting a periaxonal space width (*psw*) within the physiological range measured in the study (6.477nm) or at two extremes (*psw* = 0nm and *psw* = 20nm), and used an increasing number of segments per internode. In all scenarios, using N = 52 segments (finishing dots), ensured that a stable prediction of the conduction velocity was reached irrespective of the width of the periaxonal space.

**Table S1.** Parameters used in computational simulations of action potential conduction velocity.

| Parameter |  | Sham | Short nodes* | Alt. myelin* | iTBS* | No-Learning | Long nodes* | Alt. myelin* | Learning * |
| --- | --- | --- | --- | --- | --- | --- | --- | --- | --- |
| Internode length | $\mu\text{m}$ | 50.32 | | | | 38.89 | | | |
| Axon diameter | $\mu\text{m}$ | 0.5894 | | | | 0.5938 | | | |
| Node length | $\mu\text{m}$ | 0.8364 | 0.7735 | | 0.7735 | 0.6644 | 0.8668 | | 0.8668 |
| RMP | mV | -72 |  |  |  | -72 |  |  |  |
| Node capacitance | $\mu\text{F}/\text{cm}^2$ | 0.9 | | | | 0.9 | | | |
| Internode capacitance | $\mu\text{F}/\text{cm}^2$ | 0.9 | | | | 0.9 | | | |
| Internode leak conductance | $\text{mS}/\text{mm}^2$ | 0.1 | | | | 0.1 | | | |
| $E_L$ | mV | -84 | | | | -84 | | | |
| Axoplasmic resistivity | $\Omega\cdot\text{m}$ | 0.7 | | | | 0.7 | | | |
| Periaxonal resistivity | $\Omega\cdot\text{m}$ | 0.7 | | | | 0.7 | | | |
| Periaxonal space width | nm | 6.477 |  | 8.487 | 8.487 | 8.265 |  | 6.709 | 6.709 |
| g-ratio |  | 0.724 |  | 0.6888 | 0.6888 | 0.7009 |  | 0.7419 | 0.7419 |
| Number lamellae |  | 7 |  |  |  | 7 |  |  |  |
| Myelin membrane capacitance | $\mu\text{F}/\text{cm}^2$ | 0.9 | | | | 0.9 | | | |
| Myelin membrane conductance | $\text{mS}/\text{mm}^2$ | 1 | | | | 1 | | | |
| $g_{\text{Naf}}$ | $\text{mS}/\text{mm}^2$ | 50 | | | | 50 | | | |
| $g_{\text{Nap}}$ | $\text{mS}/\text{mm}^2$ | 0.05 | | | | 0.05 | | | |
| $g_K$ | $\text{mS}/\text{mm}^2$ | 0.8 | | | | 0.8 | | | |

\* Only detailed if different from the control (sham/no-learning) condition.

